# *Gardnerella* Diversity and Ecology in Pregnancy and Preterm Birth

**DOI:** 10.1101/2023.02.03.527032

**Authors:** Hanna L. Berman, Daniela S. Aliaga Goltsman, Megan Anderson, David A. Relman, Benjamin J. Callahan

**Affiliations:** Department of Population Health and Pathobiology, North Carolina State University, Raleigh, North Carolina; Department of Microbiology & Immunology, Stanford University School of Medicine, Stanford, California; Department of Medicine, Stanford University School of Medicine, Stanford, California; Department of Discovery, Metagenomi Inc, Emeryville, California; Infectious Diseases Section, Veterans Affairs Palo Alto Health Care System, Palo Alto, California; Bioinformatics Research Center, North Carolina State University, Raleigh, North Carolina

**Keywords:** Gardnerella, vaginal microbiome, preterm birth

## Abstract

The vaginal microbiome has been linked to numerous negative health outcomes including preterm birth. Specific taxa, including *Gardnerella* spp., have been identified as risk factors for these conditions. Historically, microbiome analysis methods have treated all *Gardnerella* spp. as one species, but the broad diversity of *Gardnerella* has recently become more apparent. In the present study, we explore the diversity of *Gardnerella* clades and genomic species in the vaginal microbiome of pregnant women and their impacts on microbiome composition and associations with preterm birth.

Shotgun metagenomic sequencing data collected longitudinally from three distinct cohorts of pregnant women were assessed. Relative abundance of *Gardnerella* clades and genomic species and other taxa was quantified, and associations between *Gardnerella* clades and signatures of the vaginal microbiome were measured. We also assessed the diversity and abundance of *Gardnerella* variants in 16S rRNA gene amplicon sequencing data from seven previously conducted studies in differing populations on the vaginal microbiome and preterm birth.

Individual microbiomes often contained multiple *Gardnerella* variants, and the number of clades was associated with increased microbial load. The genus *Gardnerella* was also associated with increased microbial load, or the ratio of non-human reads to human reads. Taxon co-occurrence patterns matched previously described community structures, and were largely consistent across *Gardnerella* clades and among cohorts. Some variants previously described as rare were prevalent in other cohorts, highlighting the importance of surveying a diverse set of populations to fully capture the diversity of *Gardnerella*.

The diversity of *Gardnerella* both across populations and within individual vaginal microbiomes has long been unappreciated, as has been the intra-species diversity of many other members of the vaginal microbiome.^1^ The broad genomic diversity of *Gardnerella* has led to its reclassification as multiple species; here we demonstrate the diversity of *Gardnerella* found within and between vaginal microbiomes. Further studies should investigate the phenotypes of *Gardnerella* variants that may underlie the mechanisms by which *Gardnerella* species may differentially shape the vaginal microbiome.

## Introduction

The microbial community that inhabits the vagina, i.e. the vaginal microbiome, both influences and reflects gynecologic and obstetric health. The vaginal microbiome plays a role in sexually transmitted infections (STIs),^2,3^ cervical cancer,^4,5^ preterm birth (PTB; delivery at <37 gestational weeks), and bacterial vaginosis (BV), a polymicrobial outgrowth of anaerobic flora.^6–8^ *Gardnerella* is a keystone genus in the vaginal microbiome.^9^ High abundance of *Gardnerella* is considered perhaps the strongest single indicator of BV^10–12^ and has been reported as one of the strongest taxonomic associations with PTB.^13,14^ *Gardnerella* is known to act in ways that can harm the host, such as producing sialidase that breaks down the protective mucins in the vagina,^15–17^ and is associated with increased host expression of inflammatory markers.^18–20^ However, over the past several decades it has become clear that *Gardnerella* is not synonymous with negative health outcomes, but is also found at varying abundances in the vagina of women without any symptoms or disease.^21^

The complexity of the relationship between *Gardnerella* and health may be explained by the phenotypic and genomic breadth of this genus that was, until recently, obscured by the classification of all *Gardnerella* as a single named species: *Gardnerella vaginalis (sensu lato*). Several studies have shown that *Gardnerella* isolates can differ in phenotypes related to pathogenicity, such as biofilm formation, adherence to epithelial cells, cytotoxicity, and interactions with *Lactobacillus* spp.^22–24^ *Gardnerella* also exhibits genomic diversity far beyond what is typically associated with a microbial species.^1,25^ Several schema have been proposed for delineating *Gardnerella vaginalis (sensu lato*) into more refined taxonomic units. Until recently, one of the most commonly used was four phylogenetic clades.^26,27^ In 2019, Vaneechoutte et al., proposed dividing *Gardnerella* into 13 species based on a 96% average nucleotide identity, and proposed names for four of these genomic species: *G. leopoldii, G. piottii, swidsinskii*, and *G. vaginalis (sensu stricto*).^28,29^ A growing number of studies have reported that *Gardnerella* clades, genomic species, or amplicon sequence variants were differently associated with specific pathogen phenotypes,^16^ BV,^21,30,31^ and PTB.^14^

High-throughput sequencing has transformed microbiome science over the past twenty years, but previous studies of the vaginal microbiome have mostly failed to distinguish between *Gardnerella* variants. Sequencing of the 16S rRNA gene has been the most common way to profile the composition of the vaginal microbiome. In these studies, it has been typical to group all *Gardnerella* together into a single taxonomic unit, precluding any possibility of detecting differences amongst variants. A few studies have used higher-resolution amplicon sequence variant (ASV) approaches to distinguish some *Gardnerella* variants based on differences in the V4 region of the 16S rRNA gene, but this yields only limited taxonomic resolution and not all V4 ASVs map uniquely to the *Gardnerella* phylogeny.^14^ Amplicon sequencing targeting other genes, such as the *cpn60* gene, has also been used.^31,32^ Some other approaches to distinguish *Gardnerella* variants within the vaginal microbiome have either yielded low resolution,^33,34^ or are based on taxon-specific marker genes that do not capture the rest of the vaginal microbiome.^25,29^ A systematic method to consistently define *Gardnerella* variants, and the ability to identify these variants in the context of the vaginal microbiome is necessary to fully understand the role of *Gardnerella* in health outcomes.

One high-throughput sequencing strategy overcomes these issues by sequencing all genomic material from all organisms in the community. This method, known as shotgun metagenomic sequencing, can distinguish between species within a genus. However, readily available computational methods for analyzing shotgun metagenome data lump together all *Gardnerella* variants. Shotgun metagenomic sequencing is still not widely used to profile the vaginal microbiome, in part due to the high proportion of host DNA in vaginal swab samples, but its use is increasing.^19,35^ In order to make the increased costs of shotgun sequencing worthwhile for studying the vaginal microbiome, new tools are needed that can make use of this richer data to accurately achieve higher resolution within important taxa such as Gardnerella.

In the present study, we establish and quantitatively validate a method to identify *Gardnerella* variants from shotgun sequencing of the vaginal microbiome. We use this method to measure the presence and abundance of six clades and 14 genomospecies of *Gardnerella* in longitudinally collected vaginal swab samples from three distinct cohorts of pregnant women. We assess the prevalence and the richness of clades and genomospecies and demonstrate and find that it is surprisingly common for all 6 clades or 14 genomospecies of *Gardnerella* to co-exist in the same vaginal microbiome. We find that presence of the genus *Gardnerella* is strongly associated with increased microbial load, and all *Gardnerella* clades were associated with increased microbial load in at least one cohort. We evaluate whether the gestational presence of different *Gardnerella* clades might differently associate with risk of preterm birth. Lastly, we assess the prevalence and abundance of amplicon sequence variants of *Gardnerella* from vaginal microbiome profiles of seven previously studied cohorts of pregnant women, and find evidence that co-colonization by multiple *Gardnerella* variants is common across a range of populations. Overall, our findings highlight the broad diversity of the *Gardnerella* genus and the rich diversity of *Gardnerella* that commonly exists within single microbiomes.

## Methods

### Gardnerella phylogeny

A core genome phylogeny of *Gardnerella* was reconstructed using complete and draft *Gardnerella* assemblies downloaded from GenBank in October 2020. The 124 assemblies were assessed for completeness and quality. First, the assemblies were verified as *Gardnerella* by aligning with blastn^36^ against the 16S ribosomal RNA blast database (downloaded from ftp://ftp.ncbi.nlm.nih.gov/blast/db/16S_ribosomal_RNA.tar.gz). Assemblies that did not align to any reference 16S sequence from any organism (five), or the best alignment by bit score was either not *Gardnerella* (two) or was *Gardnerella* but aligned with a percent identity less than 96% (one) were further assessed by aligning against the nr/nt database. Based on blast alignments with 16S rRNA and nr/nt databases, two assemblies, GCA_002871555.1 and GCA_902362445.1, were suspected to be *Lactobacillus vaginalis* and excluded.

Assemblies verified as *Gardnerella* were then assessed for completeness and quality. To assess completeness, assemblies were aligned against 40 single-copy core genes described by Mende et al (Table S1).^37^ Assemblies with 50 or more of these core genes and/or a contig N50 < 5000 were removed. One sequence was removed as a partial assembly (GCA_000263455.1) and three were removed as co-assemblies from shotgun metagenomic sequencing (GCA_902373565.1, GCA_003240925.1, GCA_003240955.1). Sequence similarity was assessed by performing all-against-all alignments with Mash^38^ version 2.2. The set of genome assemblies was then de-replicated by selecting one representative genome from each cluster of genomes within a Mash distance of 0.005 based on status as a complete genome or assembly quality based on contig N50 and L50 values.

After filtering, 85 *Gardnerella* assemblies with a mean genome length of 1.6Mbp (*SD* = 0.009 Mbp (Figure S1A) were used to reconstruct a phylogeny. Analysis of the pangenome of these 85 *Gardnerella* assemblies identified 85 core genes from a total of 12,105 genes in the pangenome (Figure S1B). The 85 *Gardnerella* assemblies were annotated with Prokka^39^ version 1.14.6 and the core genome was determined using Roary^40^ version 3.13.0 with a 95% blastp threshold.The core genes were aligned with MAFFT^41^ within Roary and used to reconstruct a maximum likelihood core-genome phylogeny with RAxML^42^ version 8.2.12 using a GTR+Gamma model. Confidence values were determined with 250 bootstrap replicates, as determined using the autoFC algorithm in RAxML.^43^ The phylogeny was subsequently rooted with *Bifidobacterium longum* 51A (NZ_CP026999.1) using the EPA algorithm in RAxML. Previously described variants, including *cpn60* variants, clades, and genomospecies are labeled on the phylogeny (Figure 1).

**Figure 1.**
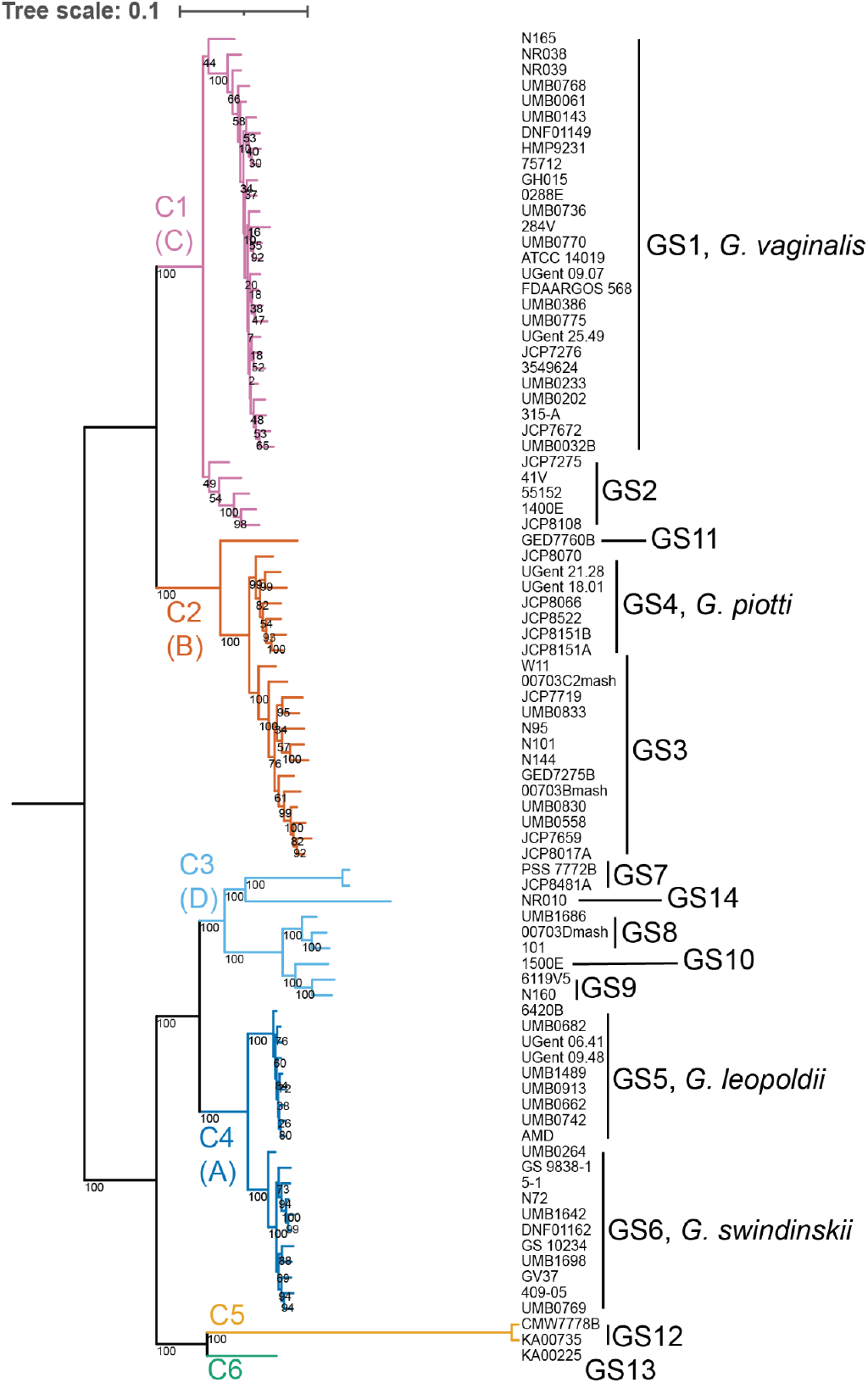
G. vaginalis core genome phylogeny: Core genome phylogeny reconstructed from 85 core genes from 85 *Gardnerella* whole genome assemblies. Core genes were determined using Roary, aligned with MAFFT, and a maximum likelihood phylogeny was reconstructed using RAxML, with a GTR+Gamma model. Confidence values on branches reflect 250 bootstrap replicates. The phylogeny was rooted with *Bifidobacterium longum* 51A, using the EPA algorithm in RAxML. Colors indicate six subspecies clades, labeled C1-C6, with previously described *cpn60* variants A through D labeled in parentheses,^27^ and lines indicate the genomospecies first described by Vaneechoutte et al.^28^

### Study Cohorts

The present study utilized shotgun metagenomic sequencing performed on vaginal swab samples from three demographically distinct cohorts of pregnant women (see Table 1). These three cohorts were from previously conducted studies in which participants were prospectively enrolled and collected vaginal swab samples were collected longitudinally through pregnancy. Shotgun metagenomic sequencing was conducted specifically for this study in two cohorts, Stanford Enriched and University of Alabama at Birmingham Enriched (UAB Enriched).^13,14^ Samples from these subjects with high relative abundance of *Gardnerella* were selected based on amplicon sequencing of the V4 region of the 16S rRNA gene. Sixty-two samples were selected from the Stanford cohort (Palo Alto, CA; 20 women; 9 of whom delivered preterm). The underlying population was predominantly white and Asian with low risk for PTB (<10%). Forty-five samples were selected from the UAB cohort (Birmingham, AL; 15 women; 10 preterm). The underlying population was predominantly African American, and women in this cohort were referred based on prior history of PTB.

**Table 1.** Cohort Demographic Information

|  | Stanford Enriched |  | UAB Enriched |  | MOMS-PI |  |  |
| --- | --- | --- | --- | --- | --- | --- | --- |
|  | Term (n=11) | Preterm (n=9) | Term (n=5) | Preterm (n=10) | Term (n=90) | Preterm (n=43) | Unknown (n=98) |
| Mean gestational age in weeks at delivery (±SD) | 39.2 (1.7) | 33.9 (2.3) | 37.8 (1.3) | 32.7 (3.4) | 40 (0.7) | 34.1 (3.4) | - (-) |
| Age |  |  |  |  |  |  |  |
| Below 18 | 0 (0%) | 0 (0%) | 0 (0%) | 0 (0%) | 2 (2%) | 1 (2%) | 1 (1%) |
| 18 to 28 | 2 (18%) | 0 (0%) | 4 (80%) | 6 (60%) | 57 (63%) | 28 (65%) | 54 (55%) |
| 29 to 38 | 4 (36%) | 6 (67%) | 1 (20%) | 3 (30%) | 27 (30%) | 12 (28%) | 39 (40%) |
| Above 38 | 0 (0%) | 0 (0%) | 0 (0%) | 0 (0%) | 4 (4%) | 2 (5%) | 2 (2%) |
| Unknown | 5 (45%) | 3 (33%) | 0 (0%) | 1 (10%) | 0 (0%) | 0 (0%) | 2 (2%) |
| Race |  |  |  |  |  |  |  |
| Asian | 2 (18%) | 0 (0%) | 0 (0%) | 0 (0%) | 0 (0%) | 0 (0%) | 2 (2%) |
| Black | 0 (0%) | 0 (0%) | 3 (60%) | 8 (80%) | 70 (78%) | 30 (70%) | 56 (57%) |
| White | 6 (55%) | 4 (44%) | 0 (0%) | 1 (10%) | 14 (16%) | 6 (14%) | 30 (31%) |
| Other | 3 (27%) | 5 (56%) | 2 (40%) | 1 (10%) | 6 (7%) | 7 (16%) | 10 (10%) |
| Ethnicity |  |  |  |  |  |  |  |
| Hispanic | 3 (27%) | 6 (67%) | 1 (20%) | 0 (0%) | 6 (7%) | 4 (9%) | 4 (4%) |
| Non-Hispanic | 5 (45%) | 2 (22%) | 4 (80%) | 9 (90%) | 84 (93%) | 39 (91%) | 93 (95%) |
| Unknown | 3 (27%) | 1 (11%) | 0 (0%) | 1 (10%) | 0 (0%) | 0 (0%) | 1 (1%) |
| Education |  |  |  |  |  |  |  |
| Less than high school | 1 (9%) | 3 (33%) | 3 (60%) | 4 (40%) | 6 (7%) | 5 (12%) | 11 (11%) |
| High school diploma or GED | 1 (9%) | 3 (33%) | 1 (20%) | 3 (30%) | 38 (42%) | 20 (47%) | 27 (28%) |
| Some college | 2 (18%) | 1 (11%) | 1 (20%) | 2 (20%) | 23 (26%) | 9 (21%) | 20 (20%) |
| Bachelor or undergraduate degree | 4 (36%) | 1 (11%) | 0 (0%) | 0 (0%) | 17 (19%) | 6 (14%) | 24 (24%) |
| Post-undergraduate degree | 2 (18%) | 1 (11%) | 0 (0%) | 0 (0%) | 4 (4%) | 3 (7%) | 14 (14%) |
| Unknown | 1 (9%) | 0 (0%) | 0 (0%) | 1 (10%) | 2 (2%) | 0 (0%) | 2 (2%) |
| Income |  |  |  |  |  |  |  |
| Under \$80,000 | 6 (55%) | 6 (67%) | 5 (100%) | 9 (90%) | 81 (90%) | 40 (93%) | 78 (80%) |
| \$80,000 or more | 4 (36%) | 1 (11%) | 0 (0%) | 0 (0%) | 4 (4%) | 1 (2%) | 15 (15%) |
| Unknown | 1 (9%) | 2 (22%) | 0 (0%) | 1 (10%) | 5 (6%) | 2 (5%) | 5 (5%) |
| Delivery Mode |  |  |  |  |  |  |  |
| Vaginal | 5 (45%) | 1 (11%) | 5 (100%) | 8 (80%) | 71 (79%) | 38 (88%) | 76 (78%) |
| Cesarean | 5 (45%) | 8 (89%) | 0 (0%) | 2 (20%) | 16 (18%) | 3 (7%) | 14 (14%) |
| Unknown | 1 (9%) | 0 (0%) | 0 (0%) | 0 (0%) | 3 (3%) | 2 (5%) | 8 (8%) |

To provide a cohort not enriched for *Gardnerella*, shotgun metagenomic sequencing data were obtained from the Multi-Omic Microbiome Study: Pregnancy Initiative (MOMS-PI)^19,44^via dbGaP (study no. 20280; accession ID phs001523.v1.p1). Sequencing data from this cohort were initially collected as part of the National Institutes of Health’s integrative Human Microbiome Project.^45^ Of the 930 samples from 243 subjects, 781 samples from 231 subjects met quality criteria (described below) and were used for analyses. A fourth cohort (MOMS-PI Enriched) was created by selecting samples from the MOMS-PI cohort that were enriched for *Gardnerella* based on subject average relative abundance of *Gardnerella* to match the distribution of subject average *Gardnerella* abundance in the Stanford Enriched and UAB Enriched Cohorts (median=47%), as determined by MetaPhlAn2 (Figure S2). This cohort included 145 samples from 42 participants. The median subject relative abundance of *Gardnerella* in the was 17% in the unenriched MOMS-PI cohort and 45% in the enriched cohort.

Due to differences in demographic and clinical characteristics (Table 1), in addition to differences in participant recruitment and sample processing protocols, all cohorts were analyzed separately.

### DNA Sequencing, Quality Control, and Filtering

#### Stanford Enriched and UAB Enriched

DNA from samples in the Stanford and UAB Enriched cohorts was extracted and sequenced using previously reported methods.^13,46^ DNA extraction was performed using the PowerSoil DNA isolation kit (Mo Bio Laboratories, Carlsbad CA, USA). Extraction was performed according to the manufacturer’s protocol except for an additional 10-minute incubation at 65°C immediately after the addition of solution C1. Barcoded TruSeq libraries were sequenced to obtain paired 150 base-pair reads on an Illumina HiSeq 2500. Alternatively, DNA was extracted with the AllPrep RNA/DNA Extraction kit (Qiagen) following the manufacturer’s instructions. DNA samples were processed at the High-Throughput Sequencing and Genotyping Unit at the University of Illinois Roy J. Carver Biotechnology Center. Specifically, shotgun genomic libraries were prepared with the Hyper Library construction kit (Kapa Biosystems) and sequenced on an Illumina HiSeq 4000 (150 bp x 2 mode).

#### ALL

Raw metagenomic reads were subjected to low quality base trimming with Sickle^47^ with parameters-n −l 100. Human reads were then filtered from samples by mapping against the human genome GRCh38.p13 with BBmap^48^ version 38.44. Samples with fewer than 1 million paired reads after Sickle filtering were excluded. Retained MOMS-PI samples contained smaller library sizes than the Stanford and UAB Enriched cohorts (Figure S3). Samples contained an average of 88%, 77%, and 91% human reads in the Stanford Enriched, UAB Enriched, and MOMS-PI samples, respectively.

### Measuring *Gardnerella* clades and genomospecies

The presence and abundance of *Gardnerella* clades and genomospecies^28^ were identified using a two-step filtering and mapping method. First, a core genome reference database was curated. The database included 70 single-copy core genes from 14 representative genome assemblies (Table S2). This set of 70 genes is a subset of the 85 core genes used to reconstruct the *Gardnerella* phylogeny and were chosen because they were single-copy. Reference assemblies were selected by first removing N165 and JCP7275 based on poor confidence in phylogenetic placement (Figure 1; bootstrap values 44 and 49). One representative core genome per genomospecies was chosen for a reference database for determining the presence and abundance of *Gardnerella* clades and genomospecies in shotgun metagenomic sequencing data. Representative core genomes were chosen based on assembly quality as complete genome status or contig N50 if there were no complete genomes in that group. If core-genomes were of equivalent quality, the core genome with the length closest to the median core-genome length (77574 bp) was chosen. Shotgun reads were then aligned against the reference database using Bowtie2^49^ version 2.1.0 and reads with a mapq score ≤ 20 were filtered and removed to retain only reads that mapped to the reference *Gardnerella* database. Reads were then binned to clades or genomospecies by a second alignment against the reference database with USEARCH^50^ version 11.0.667_i86linux32. Read identity was determined by the clade or genomic species identity of the best alignment of ≥ 99% identity.

Twelve of the 70 single-copy core genes did not resolve uniquely to a single clade and therefore alignments to these genes from all reference assemblies were not used to identify clades. Similarly, 39 of the 70 single-copy core genes did not resolve uniquely to genomospecies and reads mapping to those 39 genes were not used to identify genomospecies. Clades or genomospecies were considered present if ≥2 reads mapped to a reference gene of that variant. Validity of our mapping method to accurately identify the presence and proportion of *Gardnerella* clades and genomospecies was assessed by comparing results to the proportion of *Gardnerella* variants determined by amplicon sequencing in a subset of 28 samples on which amplicon sequencing of the V4 region of the 16S rRNA gene was performed by Pearson correlations among ASVs, clades, and genomospecies, with the ggpubr package, version 0.4.0. Additionally, we compared our *Gardnerella* mapping method to the MetaPhlAn2 results, via Pearson correlations between reads mapped to our custom database and reads mapped to *Gardnerella* references in the MetaPhlAn2 database.

### Measuring vaginal microbiome species abundance

Relative abundance of total *Gardnerella* and other taxa in the vaginal microbiome was determined using MetaPhlAn2.^51^ Microbial load for each sample was defined as the ratio of microbial reads (number of reads remaining after filtering against human reference genome with BBmap) to human reads (number of reads removed by filtering against human reference genome with BBmap). This approach of measuring microbial load by dividing the non-host reads by the host reads has been used previously in plant biology.^52^

### *Gardnerella* 16S rRNA ASVs in previously collected cohorts

To observe *Gardnerella* diversity across a wider set of cohorts, we analyzed *Gardnerella* amplicon sequencing variants (ASVs) in previous studies that profiled the vaginal microbiome in pregnancy using the V4 region of the 16S rRNA gene.^13,14,53–56^ The sequencing data were processed uniformly as part of a larger meta-analysis effort.^57^ Seven cohorts from six publications were assessed. Exact sequencing variants were resolved using DADA2^58^ version 1.12.1. Taxonomy was assigned to all ASVs in the cohorts with the assignTaxonomy function in the DADA2 package and the Silva version 138 reference database. ASVs assigned to the genus *Gardnerella* were retained. *Gardnerella* ASVs were mapped against the reference phylogeny (Figure 1C) to determine which ASVs could be resolved to clades. ASVs that were not found among any of the reference *Gardnerella* whole genome sequences were labeled “unmapped”.

### Statistical analysis

The impact of all *Gardnerella* sequence types as a group, and each of the six described clades on the ecology of the vaginal microbiome was then assessed, focusing on lactobacilli and anaerobes commonly found in the vaginal microbiome. *Lactobacillus spp*. included *Lactobacillus crispatus, L. gasseri, L. jensenii*, and *L. iners*. Anaerobes assessed included *Atopobium vaginae, Finegoldia magna, Mycoplasma hominis, Megasphaera genomospecies type 1, Megasphaera unclassified, Prevotella amnii, Prevotella bivia, Prevotella timonensis*, and *Ureaplasma parvum*. A subset of the anaerobes which are commonly discussed in the literature as important vaginal microbiome taxa are shown in the main text results, and the full list of anaerobes are in supplemental figures. Spearman’s rank correlation was performed with the ggpubr package to test the association between clades per sample and microbial load. To ensure that the association between clade counts and PTB was not due to differences in library size, we performed the analysis again after rarefying to a common microbial read depth. Human-read-filtered samples were rarefied to 100,000 reads with Seqtk version 1.3, and then the association of non-rarefied microbial load and clade counts were assessed as above. When measuring presence-absence versus microbial load or co-occurrence of taxa, presence was defined as a relative abundance greater than 0.1% were considered present. Microbial load by taxon presence absence was assessed using a Wilcoxon Rank Sum test with *p* values adjusted using the Benjamini Hochberg method in the rstatix version 0.7.0 package. Co-occurrence patterns were assessed by measuring Jaccard distance with Vegan^59^ version 2.5-7. Jaccard distance is a measure of similarity, where 1 indicates that the two taxa were not found together in samples and 0 indicates that the taxa always co-existed in samples.

To assess the impact of *Gardnerella* variants on preterm birth, mean gestational relative abundance of each clade and genomospecies in subjects who delivered at term and preterm were compared using Wilcoxon Rank Sum tests in the MOMS-PI cohort. Mean gestational clade counts and mean gestational microbial load were also compared between subjects who delivered at term and preterm using Wilcoxon Rank Sum tests in the MOMS-PI cohort. When counting the number of clades present in each sample, a clade was considered to be present if the relative abundance was greater than 0.1%. To ensure that the association between clade counts and PTB was not due to differences in library size, we performed the analysis again after rarefying using the methods described above, and then mean gestational clade counts were compared between term and preterm births

The associations between clade or genomospecies abundance and preterm birth were not assessed using Wilcoxon Rank Sum tests in the three enriched cohorts due to its impacts on *Gardnerella* abundance. Therefore, we assessed pairwise differences among the associations of each clade and genomospecies abundances and preterm birth. First, differences in mean gestational relative abundance of each pair of clades were calculated. Differences in relative abundance were transformed using a fourth-root transformation. We then assessed whether the pairwise differences varied among term and preterm births by performing logistic regression with the following model using the glm function in the R Stats package version 4.1.1:

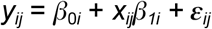

where *y_ij_* is the gestational age at delivery (term or preterm) for each subject *i* and each clade pair *j*.

We assessed *Gardnerella* ASV abundance among the seven cohorts with previously collected 16S amplicon sequencing data. We first measured the total number of *Gardnerella* ASVs across all cohorts and the number of *Gardnerella* ASVs per sample. Mean cohort abundance of *Gardnerella* ASVs was compared. Subject average relative abundance of *Gardnerella* ASVs was assessed between subjects who delivered at term and preterm using generalized linear mixed models. Subject average abundance was log10-transformed after the addition of a pseudocount of 0.00001. The following model was fit for each of the five ASVs using the lmer function in the lme4 package version 1.1.29:

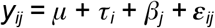

Where *y_ij_* is ASV abundance, for each subject *i* in each study *j, μ* is the population mean abundance, *τ* is the random variable for the effect of PTB on ASV abundance, and *β* is the fixed effect of each study. Study was included as a fixed effect due to the sampling, socioeconomic, racial and other differences in each study.

All analyses were conducted in the R statistical computing environment version 4.1.1^60^ using RStudio version 2022.07.1+554.^61^

## Results

### Current genomic census supports 14 genomic species and 6 distinct clades of *Gardnerella*

On October 1, 2020, we downloaded 124 *Gardnerella* genome assemblies from GenBank. After quality assessment and de-replication (see Methods), 85 whole genome *Gardnerella* assemblies were used to construct a core-genome phylogeny of *Gardnerella* (Table S2, Figure 1). Multiple schemas have been proposed for phylogenetically delineating *Gardnerella* into finer taxonomic units.^29,62^ The most commonly used are a set of four phylogenetic clades that can be differentiated by sequencing the cpn60 marker gene,^26,27^ and 13 genomic species that share <96% genomic average nucleotide identity with one another.^28^ Finally, an ad hoc classification of *Gardnerellas* into three main groups based on the sequence of the V4 region of the 16S rRNA gene has also been described.^14^ Each of these schemas for delineating *Gardnerella* are labeled on a common phylogenetic tree in Figure 1.

Our phylogeny largely recapitulates the previously described classifications of different *Gardnerellas*, but some clarifications come into focus. Our phylogeny reveals six distinct clades C1-C6. These clades reflect the four previously described clades^26^ and two additional clades comprised of isolates whose genome sequences were made available after the four initial clades were proposed in 2012. The 13 initially described *Gardnerella* genomospecies (GS1-GS13) were all clearly identifiable in our phylogeny. One additional genomospecies represented by strain NR010 was identified and labeled GS14 (see Methods). This strain was previously described as a 14th genomospecies.^29^ Species names were previously proposed for four of the genomospecies^28^: GS1, *G. vaginalis;* GS4, *G. piotii;* GS5, *G. leopoldii*, GS6, *G. swindinskii*. Here, *G. vaginalis* will be used to refer to the *G. vaginalis* GS1 genomospecies, not as per its historical usage as the entire *Gardnerella* genus. Finally, five different groups of strains were identified based on unique sequences of the V4 hypervariable region of the 16S rRNA gene (G1-G5). These correspond to the three main ASVs (G1-G3) described by Callahan et al.,^14^ with two additional ASVs (G4-G5) obtained from diverged and less-sequenced *Gardnerellas*. Of note, whereas G1, G2, G4 and G5 represent phylogenetically coherent groups of strains, G3 is highly polyphyletic (Figure 1). Going forward, “variant” will be used as a generic term for phylogenetic units including clades (C1-C6), genomospecies (GS1-GS14), and groups of strains sharing a common V4 amplicon sequence variant (G1-G5).

### Vaginal swab metagenomes from three distinct cohorts of pregnant women

Shotgun metagenomes from vaginal swab samples were obtained from three cohorts of pregnant women (see Methods).^14,19^ The first cohort -- the Multi-Omic Microbiome Study: Pregnancy Initiative (MOMS-PI; N=231 subjects, 781 samples) -- was described including publicly available shotgun sequencing data by Fettweis et al.^19^ and Serrano et al.^44^ Two further cohorts -- Stanford Enriched (N=20 subjects, 62 Samples) and University of Alabama at Birmingham Enriched (UAB Enriched; N=15 women, 45 samples) τ were previously described,^13,14^ but had their shotgun metagenomic sequencing performed for this study. Briefly, paired 150 base-pair reads were sequenced on an Illumina HiSeq 2500 platform. In these two cohorts, shotgun metagenomes were obtained from an “enriched” set of samples known to contain *Gardnerella* by presence of *Gardnerella* ASVs based on amplicon sequencing of the V4 region of the 16S rRNA gene. Therefore, the Stanford Enriched and UAB Enriched cohorts are not a random sample of the original study populations. These three cohorts were racially, ethnically, and socioeconomically distinct (Table 1). A synthetic fourth cohort (MOMS-PI Enriched; N=42 subjects, 145 samples) was generated by subsampling the MOMS-PI cohort to match the distribution of *Gardnerella* relative abundances with the Stanford Enriched and UAB Enriched cohorts (Figure S4, see Methods). All cohorts contained samples from women who delivered at term and from women who delivered preterm. Library sizes contained an average of 19.7 million reads in the Stanford Enriched cohort and an average of 19.2 million reads in the UAB Enriched cohort. The samples in these two cohorts were sequenced at a greater depth than the MOMS-PI cohort, with an average library size of 11.2 million reads (Figure S3).

A common quality control workflow was applied to data from all cohorts (see Methods). Samples in the MOMS-PI cohort contained a larger proportion of human reads (91%) than the Stanford Enriched (88%) and UAB Enriched (77%) cohorts.

### A validated method for identifying and quantifying *Gardnerella* clades and genomospecies from shotgun metagenomic sequencing

We developed a computational method to identify *Gardnerella* clades and genomospecies by mapping shotgun metagenomic sequencing reads against a custom database of Gardnerella core genes (see Methods), which resolved *Gardnerella* to clade and species levels. We validated our method by comparing results to *Gardnerella* identified by 16S rRNA gene amplicon sequencing and MetaPhlAn2. First, we compared the proportion of total *Gardnerella* belonging to each clade and genomospecies from read mapping to the proportion of each *Gardnerella* ASV (of the total *Gardnerella* ASVs) in 28 samples from the Stanford Enriched and UAB Enriched cohorts in which both shotgun metagenomic sequencing and 16S rRNA gene amplicon sequencing had been performed. Pearson correlations were used to test the association between proportion of each ASV and the sums of clade and genomospecies proportions to which those ASVs map (Figure 2A-B). Proportions of ASV G3 were not plotted or included in Pearson correlations due to its polyphyletic nature. Proportions marked by a triangle in Figure 2A-B are from the two samples in which ≥50% of *Gardnerella* amplicon sequencing reads were the G3 ASV. Summed proportions of *Gardnerella* clades and genomospecies were significantly correlated to the proportion of ASVs that map to those clades or genomospecies (*p* < 0.0001; Figure 2A-B). Proportions of clades were also significantly correlated with the sum proportions of genomospecies within each clade (*p* < 0.0001; Figure 2C).

**Figure 2:**
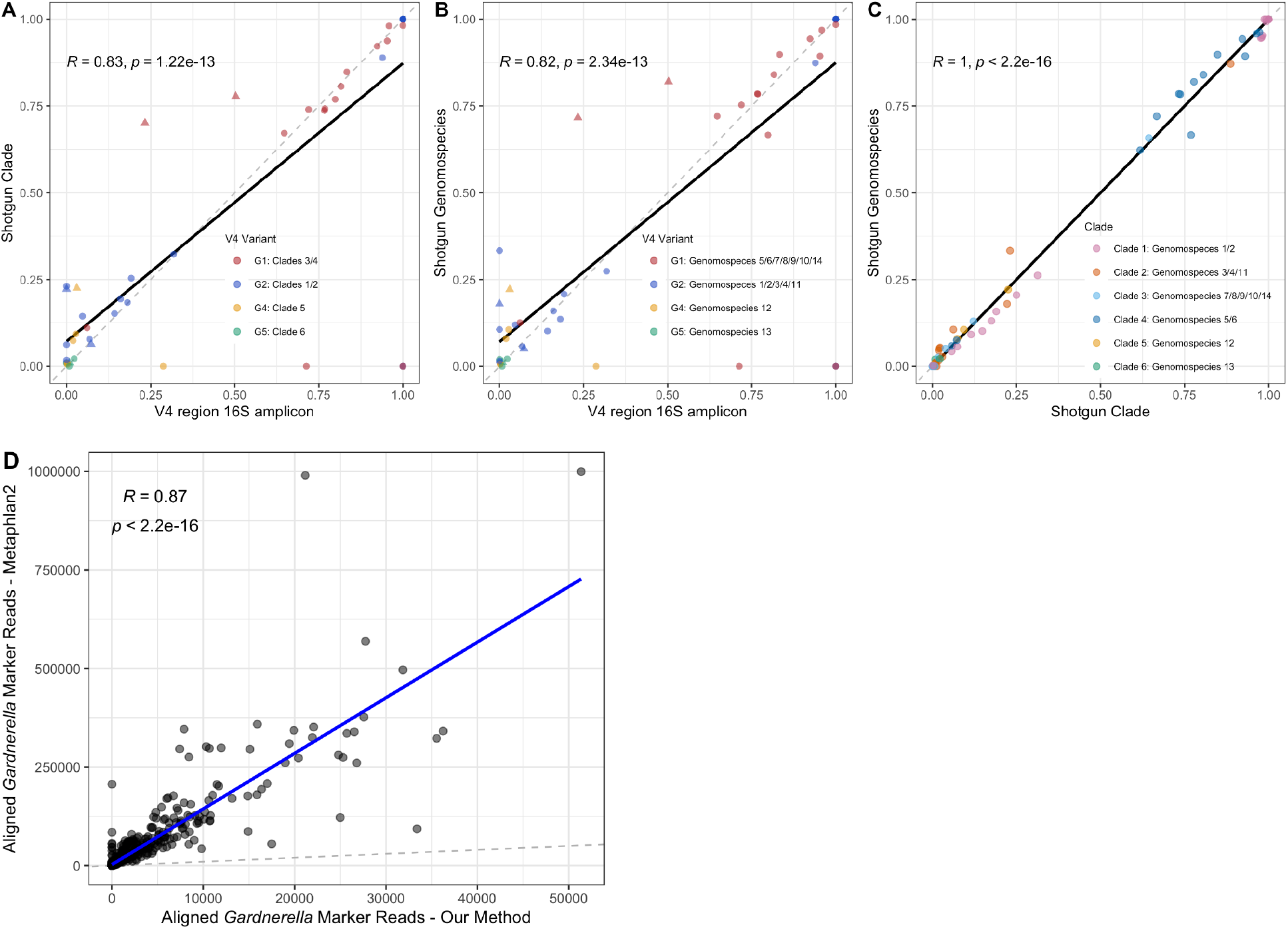
Validation of *Gardnerella* mapping method against 16S ASVs and MetaPhlan2: Pearson correlations of proportions of A) *Gardnerella* clades to V4 16S ASVs, B) *Gardnerella* genomospecies to V4 16S ASVs, and C) genomospecies to clades out of all *Gardnerella* in 28 samples from the Stanford Enriched and UAB Enriched cohorts in which both shotgun and amplicon sequencing were performed. Proportions of *Gardnerella* clades and genomospecies determined by our *Gardnerella* mapping method, and ASVs of the V4 region of the 16S region determined by amplicon sequencing. Points marked by a triangle indicate that they are from samples in which 50% or more of the *Gardnerella* reads were the G3 ASV by amplicon sequencing. Black lines fitted by OLS linear regression and gray dotted lines depict y=x D) Pearson correlation of aligned *Gardnerella* marker reads found by MetaPhlan2 vs. aligned *Gardnerella* marker reads found in samples by our mapping method. Blue line fitted by OLS linear regression, and the gray dotted line plotted as y=x.

*Gardnerella* reads measured by our mapping method correlated with *Gardnerella* quantified by MetaPhlAn2. We compared our clades and genomospecies mapping method to the results of *Gardnerella* and marker read mapping and abundance measurements determined by MetaPhlAn2. Between the Stanford Enriched and UAB Enriched cohorts, our mapping method found *Gardnerella* reads in one sample which MetaPhlAn2 labeled as *Gardnerella*-negative. One sample in which MetaPhlAn2 reported a greater than zero abundance of *Gardnerella* could not be assigned a genomospecies by our method. In the MOMS-PI cohort, where library sizes were smaller, there was more disagreement between our method and MetaPhlAn2. For example, *Gardnerella* clades were found in nine samples and *Gardnerella* genomospecies were found in two samples in which MetaPhlAn2 did not identify any *Gardnerella*. Conversely, MetaPhlAn2 found *Gardnerella* in 58 samples in which our method did not identify *Gardnerella* clades and 127 samples in which our method did not identify genomospecies. However, we found that overall, the number of *Gardnerella* reads mapped to our database strongly correlated with the number of *Gardnerella* reads mapped to the *Gardnerella* reference marker genes in the MetaPhlAn2 database (Figure 2D).

### Many variants of *Gardnerella* often coexist in a single microbiome

When Gardnerella was detected in a vaginal swab sample, multiple coexisting variants were often present (Figure 3). In every cohort, vaginal swab metagenomes in which *Gardnerella* was present typically contained more than one clade (mean=2.93) and more than one genomospecies (mean=3.95). The greatest number of clades (mean=4.78) and genomospecies (mean=9.18) per sample were found in the UAB Enriched cohort (Figure 3A-B), and the number of clades (mean=3.77) and genomospecies (mean=5.38) was also higher in the MOMS-PI Enriched than in the broader MOMS-PI cohort (mean clades=2.66; mean genomospecies=3.18). The richness of *Gardnerella* variants in some samples was striking. In each cohort at least one sample contained all 14 *Gardnerella* genomospecies, and 26.67% of samples in the UAB Enriched cohort contained every genomospecies.

**Figure 3.**
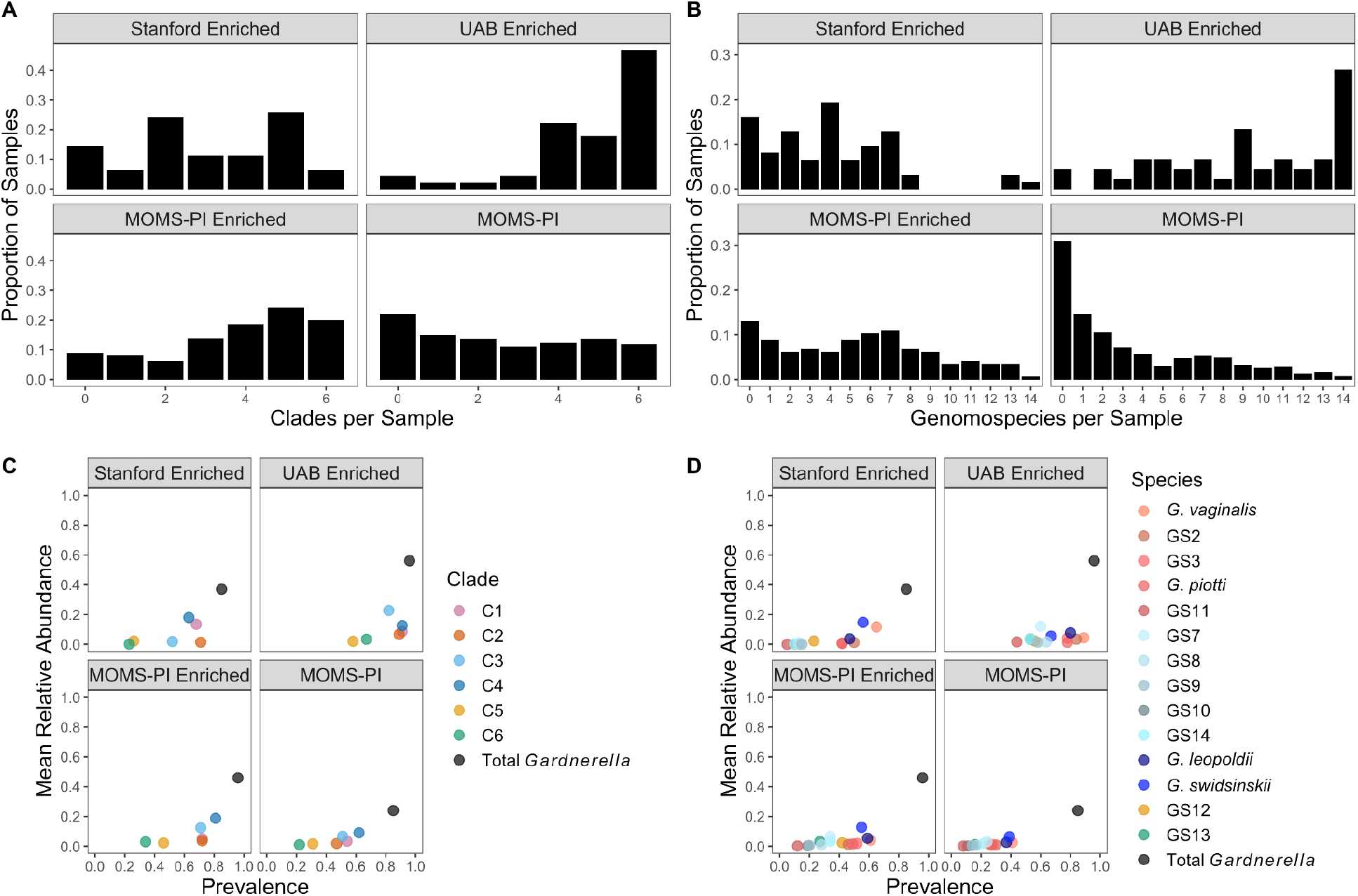
Prevalence and abundance of *Gardnerella* variants across the four cohorts: Histograms of A) clades and B) genomospecies per sample in each cohort. C) Mean relative abundance by prevalence of total *Gardnerella* and *Gardnerella* clades or D) genomospecies. Uncharacterized (U.) *Gardnerella* reflects samples where *Gardnerella* was found by MetaPhlAn2 but clades were not detected by *Gardnerella* mapping method. Abbreviations: C1 = *Gardnerella* clade 1, C2 = *Gardnerella* clade 2, C3 = *Gardnerella* clade 3, C4 = *Gardnerella* clade 4, C5 = *Gardnerella* clade 5, C6 = *Gardnerella* clade 6.

Consistent patterns in the abundance and prevalence of different *Gardnerella* variants across cohorts were evident, but specific examples show that meaningful differences exist between populations. Clades 1, 2, 3 and 4 were more prevalent and abundant than clades 5 and 6 in all cohorts (Figure 3C). However, clade 3, which has previously been described as rare based on a predominantly white cohort,^31^ was observed in high abundance and prevalence in the predominantly African-American cohorts described here, including appearing in over half of all samples from the unenriched MOMS-PI cohort. Clades 5 and 6 were the least prevalent clades in all cohorts except for UAB Enriched, where they were present in 58% and 67% of all samples, respectively. All clades and genomospecies were more prevalent and more abundant in the UAB Enriched cohort than in any other cohort, and enriching for *Gardnerella* in the MOMS-PI cohort increased the prevalence and abundance of most clades and genomospecies (Figure 3C-D).

### Presence and diversity of *Gardnerella* is associated with increased microbial load

Shotgun metagenomic sequencing samples from all the DNA in a sample, including DNA from host sources such as the vaginal epithelial cells that are collected along with the target microbiota on a vaginal swab. Here we used this property to define a proxy for the “microbial load” of the vaginal microbiome as the ratio of microbial reads to human reads in shotgun sequencing of vaginal swabs. One way to define the microbial load on an epithelial surface, such as the vaginal microbiome, is the number of microbes per unit of surface area (e.g., microbes per mm^2). Swabs do not collect a standardized amount of sample and thus swab-to-swab variation in the layers of epithelium were disturbed and thus the amount of host cells collected on the swab can add noise to our proxy measurement. Nevertheless, this noise should be uncorrelated with the true microbial load, and so this proxy measurement should still be effective at discerning patterns with sufficiently large effect sizes.

Systematic differences in microbial load were observed between cohorts, and as a function of the presence and diversity of *Gardnerella* in the vaginal microbiome. Vaginal swab samples from the UAB Enriched and MOMS-PI Enriched cohorts had greater microbial loads than did those from the Stanford Enriched and MOMS-PI cohorts (Figure S5). Increasing diversity of *Gardnerella* in the vaginal microbiome was significantly associated with increased microbial load. In every cohort, as the number of clades detected in a sample increased, so did the average microbial load (Figure 4A). To ensure that this was not a result of greater sequencing depth of microbial reads, clades per sample were measured after rarefying to a common microbial read depth of 100,000 reads (Figure S6). Clades per sample in rarefied samples were associated with greater microbial load, which was measured from non-rarefied samples only. Microbial load coul Samples in which *Gardnerella* (Total *Gardnerella* or TG) was present had a higher microbial load than samples without *Gardnerella*, in all cohorts by Wilcoxon Rank Sum (Figure 4B). All clades were significantly associated with increased microbial load in at least one cohort, and clades 1, 3, and 4 were significantly associated with increased microbial load in all cohorts*. L. crispatus* was associated with decreased microbial load, but *L. iners* was not. Presence of *A. vaginae*, another anaerobe associated with BV, was also associated with increased microbial load in all cohorts. Of the additional anaerobes assessed, unclassified *Megasphera* were significantly associated with greater microbial load in all cohorts (Figure S7).

**Figure 4.**
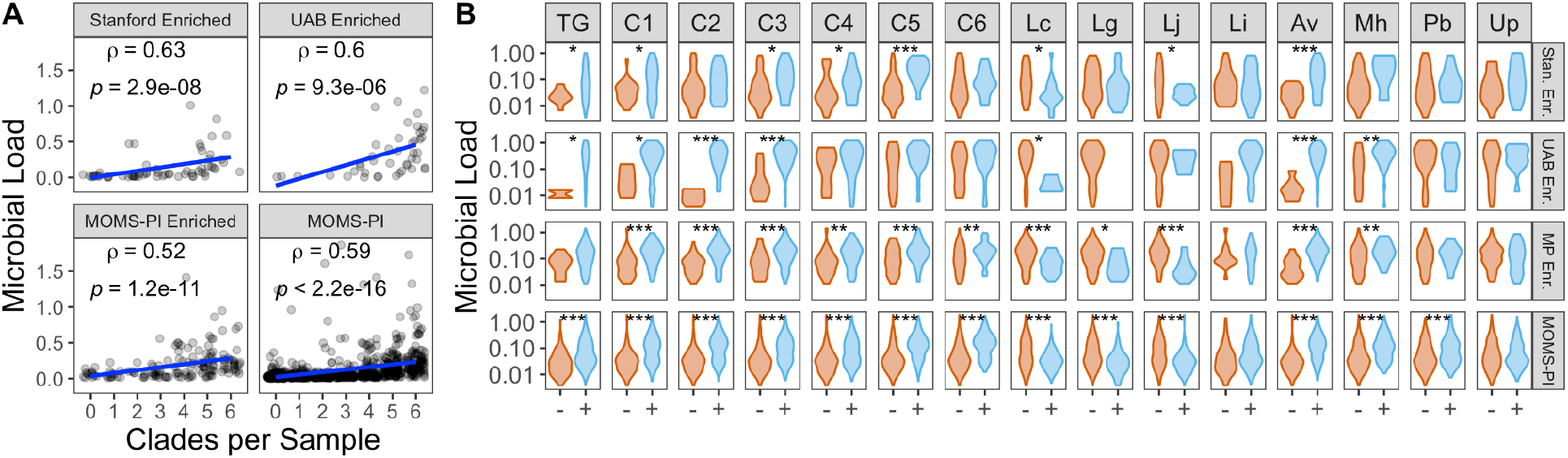
*Gardnerella* and increases in microbial load: A) Microbial load by number of clades per sample. Microbial load was significantly associated with clades per sample in all four cohorts by Spearman’s Rank correlation. Unadjusted *p* values shown. B) Microbial load by presence/absence of *Gardnerella* clades and other species in the vaginal microbiome. *p* values reflect one-sided Wilcoxon Rank Sum tests with *p* values adjusted using the Benjamini Hochberg method. *: *p* < 0.05; **: *p* < 0.01; ***: *p* < 0.00. Direction of Wilcoxon Rank Sum test determined by previous associations with microbial load: *Gardnerella* and other anaerobes associated with increased microbial load and *Lactobacillus* spp. associated with decreased microbial load. Abbreviations: TG = Total *Gardnerella*, C1 = *Gardnerella* clade 1, C2 = *Gardnerella* clade 2, C3 = *Gardnerella* clade 3, C4 = *Gardnerella* clade 4, C5 = *Gardnerella* clade 5, C6 = *Gardnerella* clade 6, Lc = *Lactobacillus crispatus*, Av = *Atopobium vaginae*, Fm = *Finegoldia magna*, Mh = *Mycoplasma hominis*, Pb = *Prevotella bivia*, Up = *Ureaplasma parvum*. Full list of taxa tested in Figure S6.

### Patterns of co-existence between *Gardnerella* and other vaginal microbes are largely consistent across clades

We investigated the relationship between total *Gardnerella* and specific *Gardnerella* clades with the rest of the vaginal microbiome, focusing on several other key taxa including the main four vaginal *Lactobacillus* species. As expected, the Enriched cohorts have greater average relative abundances of *Gardnerella* as subjects in these cohorts were selected for having detectable *Gardnerella* by 16S rRNA gene sequencing. Other notable patterns across cohorts were the higher prevalence and abundance of *L. crispatus* and *L. gasseri* in the predominantly white Stanford Enriched cohort, and higher prevalence and abundance of *L. iners* in the predominantly African-American UAB Enriched and MOMS-PI cohorts (Figure 5A). These patterns also appeared in the original Stanford and UAB cohorts based on 16S amplicon sequencing.^14^ The co-occurrence of *Gardnerella* and other taxa in the vaginal microbiome was measured using Jaccard distance (Figure 5B). The most common *Gardnerella* clades 1 through 4 typically co-occurred with each other and with *A. vaginae* and *L. iners*, but not with other *Lactobacillus* spp., although these patterns were weakest in the Stanford Enriched cohort. The less prevalent *Gardnerella* clades 5 and 6 appeared to co-occur with more common clades less frequently than the common *Gardnerella* clades. Clade 6 differed the most from the common clades in terms of its associations with other vaginal taxa, in particular not co-occurring with *L. iners* or *A. vaginae* nearly as strongly if at all. These co-occurrence patterns were largely consistent among all cohorts, including both enriched and unenriched cohorts. Co-occurrence patterns among the full list of anaerobes tested confirmed the patterns observed (Figure S8).

**Figure 5.**
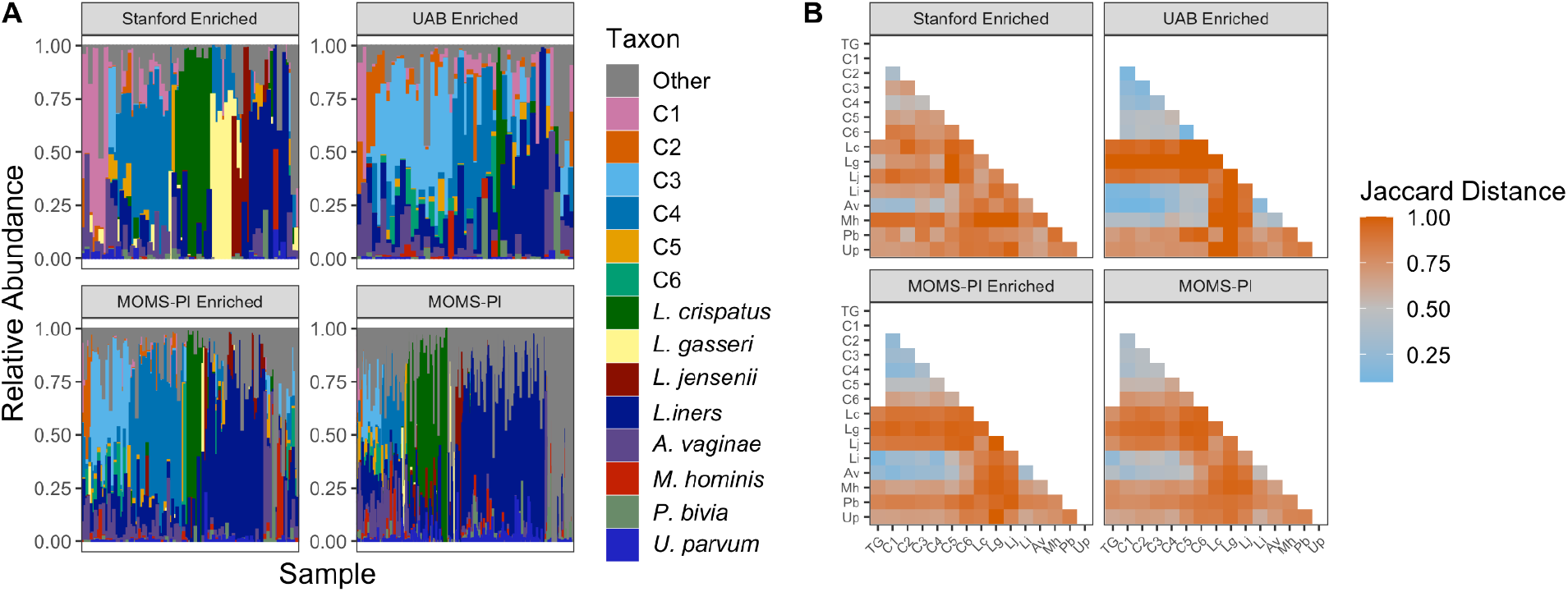
*Gardnerella* impacts on signatures of the vaginal microbiome: A) Relative abundances of species in the vaginal microbiome and *Gardnerella* clades in each sample. B) Heat map depicting Jaccard distances among common taxa in the vaginal microbiome. Abbreviations: TG = Total *Gardnerella*, C1 = *Gardnerella* clade 1, C2 = *Gardnerella* clade 2, C3 = *Gardnerella* clade 3, C4 = *Gardnerella* clade 4, C5 = *Gardnerella* clade 5, C6 = *Gardnerella* clade 6, Lc = *Lactobacillus crispatus*, Av = *Atopobium vaginae*, Fm = *Finegoldia magna*, Mh = *Mycoplasma hominis*, Pb = *Prevotella bivia*, Up = *Ureaplasma parvum*. Additional anaerobes assessed shown in Figure S8.

### Increased *Gardnerella* richness and microbial load are associated with PTB

The number of *Gardnerella* clades detected in gestational vaginal swab samples and the measured microbial load were both associated with preterm birth (PTB) outcomes in the MOMS-PI cohort (Figure 6C-D). We tested associations with PTB only in the MOMS-PI cohort, because the enrichment procedure that selected samples for metagenomic sequencing based on prior knowledge of *Gardnerella* presence could interfere with standard testing assumptions when assessing associations with *Gardnerella*. Increased number of Gardnerella clades per sample and increased microbial load were significantly associated with PTB by Wilcoxon Rank Sum test (*p* = 0.039, *p* < 0.01, respectively). To ensure that the association between mean gestational clade count and PTB was not a function of library size, clade counts were also assessed after rarefying to a common microbial read depth (see Methods) but were not associated with PTB (*p* = 0.057; Figure S9).

**Figure 6.**
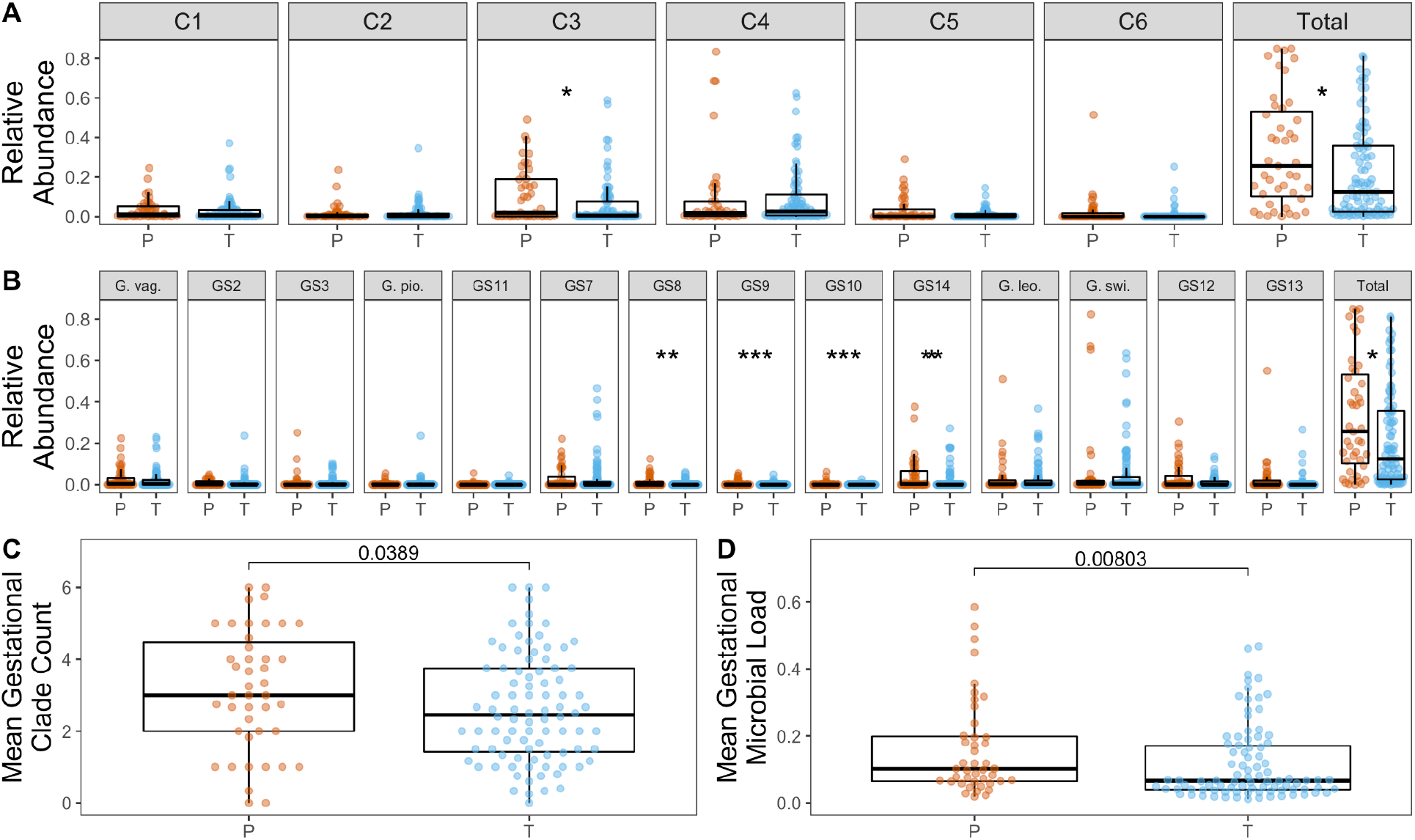
*Gardnerella* and preterm delivery status: Subject average relative abundance of *Gardnerella* and specific A) clades and B) genomospecies by preterm delivery status (Preterm birth is defined as delivery at < 37 weeks). C) Mean number of clades per subject by preterm delivery status. D) Mean bacterial load per subject by preterm delivery status. All *p* values reflect one-directional Wilcoxon Rank Sum tests. *: *p* < 0.05; **: *p* < 0.01; ***: *p* < 0.001. Note, *p* values not adjusted for multiple comparisons.

No clear differences between the associations of different *Gardnerella* variants and PTB were identified. Mean gestational relative abundance of total *Gardnerella* and clade 3 was significantly associated with PTB in the MOMS-PI cohort by Wilcoxon Rank Sum test (unadjusted *p* = 0.05, *p* = 0.01, respectively; Figure 6A). Similarly, four genomospecies within clade 3, *Gardnerella* sp. 8, *Gardnerella* sp. 9, *Gardnerella* sp. 10, and *Gardnerella* sp. 14, were significantly associated with PTB by Wilcoxon Rank Sum test (unadjusted *p* = 0.002, *p* < 0.001, *p* < 0.001, *p* = 0.01 respectively; Figure 6B). The fifth species in clade 3, *Gardnerella* sp. 7, was not significantly associated with PTB. However, when measuring pairwise differences in the associations of each clade and preterm birth in all four cohorts using logistic regression, no significant differences were found (Figure S10), perhaps because of insufficient power.

### Survey of amplicon sequencing studies finds broad diversity of *Gardnerella* and high *Gardnerella* richness across varying sample populations

In order to survey a broader set of populations, but with a more limited intra-*Gardnerella* resolution, we leveraged 16S rRNA gene amplicon sequencing data from previously conducted studies on the vaginal microbiome in pregnancy. We considered six previous studies that profiled the vaginal microbiome during pregnancy using amplicon sequencing of the V4 region of the 16S rRNA gene (Table S3). These cohorts were sampled from diverse populations and geographic regions including multiple regions of the United States and one cohort each from Peru and Canada.

A common bioinformatics workflow was used to define ASVs in each of these studies, yielding a total of 85 unique *Gardnerella* ASVs across all samples and all cohorts.^57^ However, many of these variants were found in very few samples and at very low abundances (Figure S12). 99% of all Gardnerella reads were contained in just the top 5 ASVs, which correspond to G1-G5 described above. These top five ASVs were found in all cohorts. The number of unique *Gardnerella* ASVs per sample varied among cohorts, but samples frequently contained more than one *Gardnerella* ASV, up to a maximum of eight ASVs (Figure 7A). A greater number of ASVs per sample was common in the UAB cohort of Callahan et al., 2017, consistent with our clade- and genomospecies-level observations in this cohort using shotgun metagenomic data. Mean relative abundance of the five most common variants, which are also found in our reference phylogeny, in each cohort is shown in Figure 7B.

**Figure 7.**
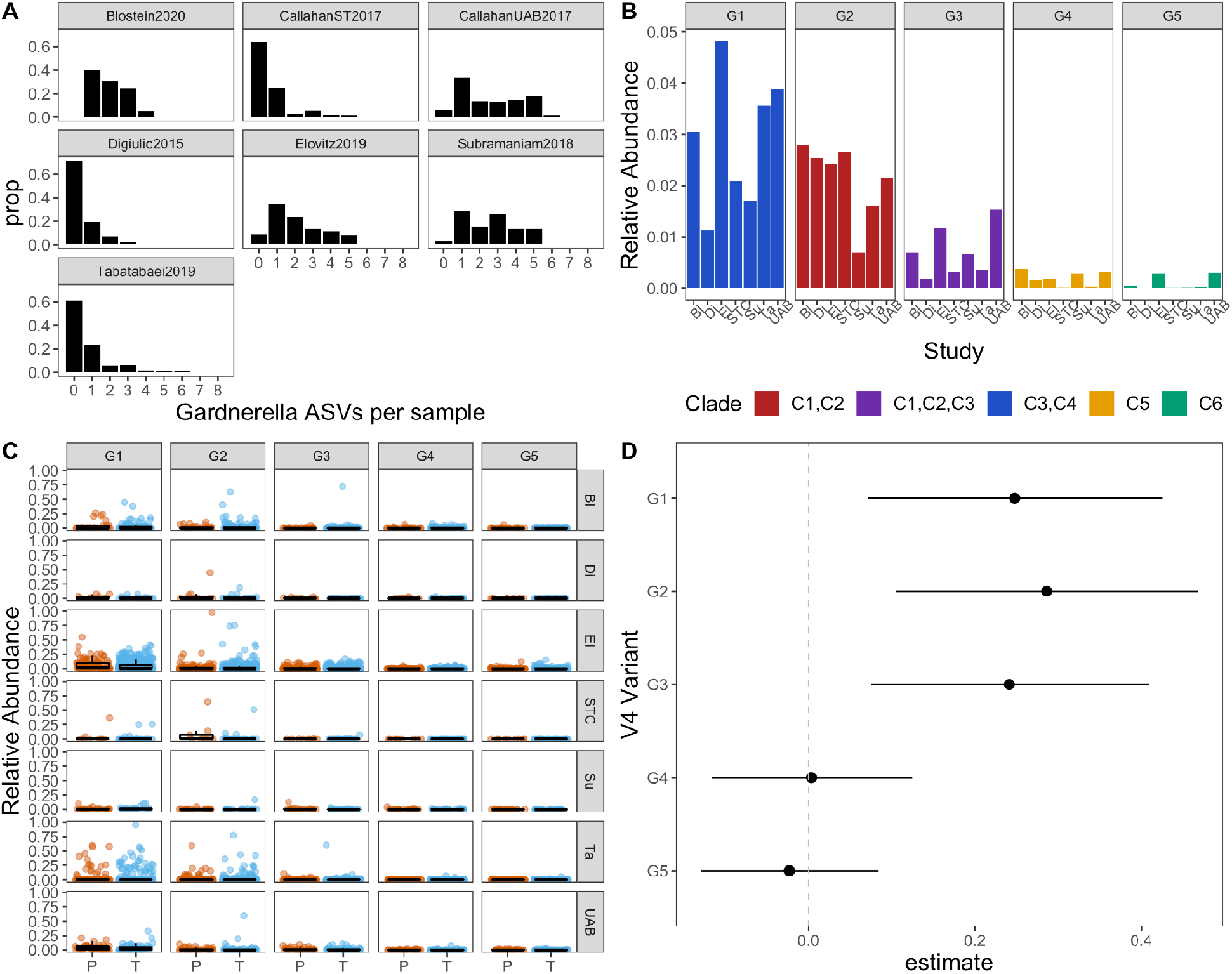
*Gardnerella* ASVs in samples from 7 cohorts profiling the vaginal microbiome with amplicon sequencing of the V4 region of the 16S rRNA gene during pregnancy: A) *Gardnerella* ASVs per sample. B) Mean relative abundance of *Gardnerella* ASVs in each cohort colored by clade. “Unmapped” indicates ASVs that were not mapped to reference *Gardnerella* phylogeny. C) Mean relative abundance of *Gardnerella* ASVs per subject in subjects that delivered (P)reterm or at (T)erm. D) Estimates and 95% confidence intervals from generalized linear mixed models of V4 variant abundance as a function of preterm delivery with study as a random variable.

We measured the association of subject average relative abundance of the top 5 ASVs with PTB (Figure 7C) using generalized linear mixed models (see Methods). A model was fitted to measure the association of PTB with log_10_ subject average relative abundance of each of the five ASVs, with study included as a random variable. Coefficient estimates and 95% confidence intervals suggested that variants G1, G2, and G3, which map to clades 1 through 4 were associated with PTB, but variants G4 and G5 (clades 5 and 6) were not (Figure 7D).

## Discussion

The present study established and validated a method to resolve *Gardnerella* to clades and genomic species in shotgun metagenomic sequencing data and assessed the diversity and ecology of *Gardnerella* variants in the vaginal microbiome in pregnancy and preterm birth. It is immediately apparent that *Gardnerella* is a diverse and cosmopolitan genus, with multiple species (and likely strains) often existing in a single microbiome and diversity shared across globally disparate populations.

The present study demonstrated the utility of shotgun metagenomic sequencing data for resolving *Gardnerella* below the genus level; however, this technology remains costly, especially in low biomass environments like the vaginal microbiome, where most of the sequences will be from the host. A more cost-effective strategy may be amplicon sequencing with a marker gene that can discriminate between *Gardnerella* species. However, relying on a single amplicon sequence can produce ambiguous placements, or placements that differ from those using genome-wide average nucleotide identity. For example, the *cpn60* sequence has been proposed as a way to discriminate *Gardnerella* species,^31^ but this may not be suitable for discriminating between phylogenetically similar species. The *cpn60* sequence of isolate N160 (GenBank Accession GCA_003408775.1) is most similar to the genomospecies 10 isolate 1500E, but would be considered genomospecies 9 by average nucleotide identity, with an average nucleotide identity of about 98% with isolate 6119V5 (see Supplemental Information). Isolate N160 was also placed in genomospecies 9 in our core genome phylogeny. Further investigation may provide additional marker candidates that better resolve *Gardnerella* species.

Our method to identify *Gardnerella* clades and genomospecies using shotgun metagenomic sequencing provided further evidence that multiple *Gardnerella* variants often co-exist in the vaginal microbiome.^25,29,31^ Our assessment of previously collected V4 16S rRNA gene amplicon sequencing data from seven cohorts also demonstrated that the number of *Gardnerella* variants in a single vaginal microbiome may vary across populations. This may have clinical relevance, as the presence of multiple *Gardnerella* variants has been associated with BV.^21,63^ Therefore, the ability to simultaneously measure multiple *Gardnerella* variants within the vaginal microbiome is important for understanding the ecology of the vaginal microbiome and potential impacts on human health.

Our results emphasized the diversity of *Gardnerella* and highlighted the importance of adequate sampling from varied cohorts. Six clades and 14 genomic species of *Gardnerella* were detected in three distinct cohorts using shotgun metagenomic sequencing. Among the seven cohorts previously profiled using amplicon sequencing, 85 *Gardnerella* ASVs of the V4 region of the 16S rRNA gene were identified. Only five of these 85 ASVs were able to be mapped to reference *Gardnerella* whole genome sequences. Therefore, the currently available whole-genome sequences of *Gardnerella* may represent a small fraction of the true diversity of *Gardnerella*. The best characterized clades (1 through 4) were the most frequent among all cohorts assessed, and some clades were found at greater abundances than previously described. For example, clade 3 has previously been described as rare;^64^ however, this variant was the most abundant clade in the UAB cohort and was also present in more than 50% of samples in our three shotgun cohorts. Additionally, the abundance of less common variants, such as clades 5 and 6, varied widely across studies and populations.

The co-occurrence of multiple *Gardnerella* strains in a single vaginal microbiome was found among all cohorts, and has been described in multiple studies. *Gardnerella* species may occupy separate ecological niches in order to exist simultaneously. In one *in vitro* assessment, *Gardnerella* isolates from clade 3 were found to be generalists, able to use significantly more carbon sources than isolates from other clades.^64^ The authors hypothesized that this ability to utilize separate carbon sources would allow clade 3 *Gardnerella* species to persist at lower relative abundances in the vaginal microbiome in the presence of other *Gardnerella* species. Additionally, co-culture experiments have demonstrated that the presence of other *Gardnerella* clades can increase the growth of clade 3 *Gardnerella* isolates, but suppresses the growth of other clades.^65^ Cell-free supernatants had no effect on growth, suggesting that these interactions are contact-dependent. Spatial separation could be important to maintain growth of other *Gardnerella* clades. This has not been described in the vaginal microbiome but has been described in other human-associated microbiomes such as the tongue.^66^

We did not find strong support for differences in the impacts of *Gardnerella* variants on signatures of the microbial community. In addition to measuring relative abundance, we also were able to obtain a measure of microbial load. Similar methods have been employed using shotgun metagenomic sequencing data from other fields.^52,67^ Validity of this method was supported by previously established relationships between taxa and microbial load. For example, *Lactobacillus crispatus*, which is associated with vaginal health, was associated with a decreased microbial load. *Gardnerella* and *Atopobium vaginae*, which are both implicated in the etiology of BV,^10^ were associated with increased microbial load. All clades were associated with microbial load in at least one cohort. *Gardnerella* was strongly associated with increased microbial load, consistent with its long-standing association with BV. The presence of multiple clades was associated with increased microbial load, suggesting that *Gardnerella* richness may be implicated in negative health outcomes.

Previous studies have demonstrated differences in *Gardnerella* co-occurrence^31^ and *in vitro* competitive growth patterns.^65^ In the present study, *Gardnerella* clades often co-occurred, and *Gardnerella* and individual clades appeared to exclude *L. crispatus*, *L. jensenii*, and *L. gasseri*, consistent with previous findings that these lactobacilli often dominate the vaginal microbiome^9^ and prohibit growth of anaerobes by maintaining a low pH. Contrastingly, *L. iners* co-occurred with *Gardnerella* and individual clades in the UAB Enriched and both MOMS-PI cohorts, a pattern that has been previously demonstrated.^9,13^ *Gardnerella* clades co-occurred with *A. vaginae* across all cohorts. As noted above, *Gardnerella* and *A*. *vaginae* are both implicated in the etiology of BV,^10^ and *in vitro* experiments suggest that *Gardnerella* may be important for growth and viability of *A. vaginae.^68^* We did not find major differences among clades, but the use of enriched cohorts may prevent the ability to extrapolate these findings to broader populations. However, similar trends were found in the unenriched MOMS-PI cohort.

Although we were able to observe the prevalence and abundance of *Gardnerella* clades and species and some relationships with signatures of vaginal microbiome composition, we were unable to describe the specific dynamics among *Gardnerella* clades or genomic species. For example, it is unknown whether certain clades act as pioneers that create a suitable environment for other variants to grow, or whether multiple clades shape the microbiome together. A proposed etiology of BV suggests that virulent *Gardnerella* strains initiate the BV biofilm along with *P. bivia* and *A. vaginae*. Additionally, it is unknown how *Gardnerella* variants are initially colonizing the vaginal microbiome, although there is evidence that *Gardnerella* is sexually transmitted.^63,69^

In the MOMS-PI cohort, in which samples were not enriched for *Gardnerella*, we found that clade 3 and four of the five genomospecies within this clade were associated with preterm birth. Previous studies have assessed the association between *Gardnerella* variants and BV or preterm birth using different methods and classifying schema. Clades 1 through 4 have been associated with BV in at least one of these studies, and clades 5 and 6 were typically not measured.^21,30,32,70^ Some studies have evaluated the associations of these genomic species with BV, and have also found varying results.^31,71^ We also found that the number of *Gardnerella* clades present was associated with PTB (in non-rarefied samples), and presence of multiple clades has been associated with BV.^21,70^ Fewer studies have included PTB as the outcome variable than BV, and results have also been mixed. A study that included a larger sampling of the Stanford and UAB Enriched cohorts found that a V4 variant of the 16S rRNA gene which maps to clades 1 and 2 was associated with PTB in its Stanford cohort, but not its UAB cohort.^14^ Another study did not find any variants associated with PTB.^34^ Often these studies contain low sample sizes and may be underpowered. We also failed to find any differences among the associations of clades or genomospecies with PTB. Therefore, these conflicting results demonstrate the necessity for additional well-powered studies in multiple populations with standardized methods for sampling, measuring, and classifying *Gardnerella* variants.

Future studies may address potential functional differences in the gene content and expression of *Gardnerella* variants in the vaginal microbiome. Additional studies may also investigate the mechanisms by which various *Gardnerella* variants may differentially interact with other taxa in the vaginal microbiome and shape health outcomes. The present study adds to the growing evidence of vast diversity within the genus *Gardnerella*, and additional surveying of *Gardnerella* in varying populations will be necessary to fully understand the diversity of *Gardnerella* in the vaginal microbiome and its association with human health.

## Author Contributions

BJC and DAR conceived of the study. BJC and DSAG selected samples for shotgun metagenomic sequencing. DSAG processed samples and performed sequence dataquality control in the Stanford Enriched and UAB Enriched cohorts. HLB performed analyses under supervision of BJC. MA organized amplicon sequence data and associated metadata and assisted with related analyses under the supervision of HLB and BJC. HLB drafted the manuscript under supervision of BJC and all authors approved of the final manuscript.

## Acknowledgements

We thank Caizhi Huang for providing the amplicon sequence data and associated metadata used in this study. We thank Gregory A. Buck, Ph.D. for providing additional metadata for the MOMS-PI cohort. We thank Alvaro Hernandez, Ph.D. and Chris Wright at the University of Illinois Roy J. Carver Biotechnology Center for guidance with library preparation and sequencing.

## Ethics Statement

The study was approved by the Institutional Review Board of North Carolina State University (Protocol Number 23575) and by the Institutional Review Board at Stanford University (Protocol Number 21956).

## Supplemental Information

Code for analyses and knit R markdowns producing figures available at https://github.com/hannalberman/pregnancy_metagenome

## Supplementary Tables

**Table S1.** Single-copy core genes for completeness testing

| Gene | Protein | Protein ID | Locus Tag | Species |
| --- | --- | --- | --- | --- |
| ychF | GTP-binding protein YchF | ADP38787.1 | HMPREF0421_20705 | Gardnerella vaginalis 14019 |
| pheS | phenylalanine-tRNA ligase, alpha subunit | ADP38666.1 | HMPREF0421_20584 | Gardnerella vaginalis 14019 |
| argS | arginine--tRNA ligase | ADP39400.1 | HMPREF0421_21318 | Gardnerella vaginalis 14019 |
| rpsL | ribosomal protein S12 | ADP38559.1 | HMPREF0421_20477 | Gardnerella vaginalis 14019 |
| rpsG | ribosomal protein S7 | ADP38560.1 | HMPREF0421_20478 | Gardnerella vaginalis 14019 |
| rpsB | ribosomal protein S2 | ADP38889.1 | HMPREF0421_20807 | Gardnerella vaginalis 14019 |
| rplK | ribosomal protein L11 | ADP38441.1 | HMPREF0421_20359 | Gardnerella vaginalis 14019 |
| rplA | ribosomal protein L1 | ADP38442.1 | HMPREF0421_20360 | Gardnerella vaginalis 14019 |
| rpoC | DNA-directed RNA polymerase, beta' subunit | ADP39290.1 | HMPREF0421_21208 | Gardnerella vaginalis 14019 |
| rplC | 50S ribosomal protein L3 | ADP39365.1 | HMPREF0421_21283 | Gardnerella vaginalis 14019 |
| rplD | 50S ribosomal protein L4 | ADP39364.1 | HMPREF0421_21282 | Gardnerella vaginalis 14019 |
| rplB | ribosomal protein L2 | ADP39362.1 | HMPREF0421_21280 | Gardnerella vaginalis 14019 |
| rplV | ribosomal protein L22 | ADP39360.1 | HMPREF0421_21278 | Gardnerella vaginalis 14019 |
| rpsC | ribosomal protein S3 | ADP39359.1 | HMPREF0421_21277 | Gardnerella vaginalis 14019 |
| rplN | ribosomal protein L14 | ADP39355.1 | HMPREF0421_21273 | Gardnerella vaginalis 14019 |
| rplE | ribosomal protein L5 | ADP39353.1 | HMPREF0421_21271 | Gardnerella vaginalis 14019 |
| rpsH | ribosomal protein S8 | ADP39351.1 | HMPREF0421_21269 | Gardnerella vaginalis 14019 |
| rplF | ribosomal protein L6 | ADP39350.1 | HMPREF0421_21268 | Gardnerella vaginalis 14019 |
| rpsE | ribosomal protein S5 | ADP39348.1 | HMPREF0421_21266 | Gardnerella vaginalis 14019 |
| rpsM | 30S ribosomal protein S13 | ADP39342.1 | HMPREF0421_21260 | Gardnerella vaginalis 14019 |
| rpsK | 30S ribosomal protein S11 | ADP39341.1 | HMPREF0421_21259 | Gardnerella vaginalis 14019 |
| rplM | ribosomal protein L13 | ADP39369.1 | HMPREF0421_21287 | Gardnerella vaginalis 14019 |
| rpsI | ribosomal protein S9 | ADP39368.1 | HMPREF0421_21286 | Gardnerella vaginalis 14019 |
| hisS | histidine--tRNA ligase | ADP38915.1 | HMPREF0421_20833 | Gardnerella vaginalis 14019 |
| serS | serine--tRNA ligase | ADP38497.1 | HMPREF0421_20415 | Gardnerella vaginalis 14019 |
| rpsO | ribosomal protein S15 | ADP38416.1 | HMPREF0421_20334 | Gardnerella vaginalis 14019 |
| rpsS | ribosomal protein S19 | ADP39361.1 | HMPREF0421_21279 | Gardnerella vaginalis 14019 |
| rpsQ | 30S ribosomal protein S17 | ADP39356.1 | HMPREF0421_21274 | Gardnerella vaginalis 14019 |
| rplP | ribosomal protein L16 | ADP39358.1 | HMPREF0421_21276 | Gardnerella vaginalis 14019 |
| rplO | ribosomal protein L15 | ADP39346.1 | HMPREF0421_21264 | Gardnerella vaginalis 14019 |
| secY | preprotein translocase, SecY subunit | ADP39345.1 | HMPREF0421_21263 | Gardnerella vaginalis 14019 |
| rpoA | DNA-directed RNA polymerase, alpha subunit | ADP39340.1 | HMPREF0421_21258 | Gardnerella vaginalis 14019 |
| cysS | cysteine--tRNA ligase | ADP39220.1 | HMPREF0421_21138 | Gardnerella vaginalis 14019 |
| rplR | ribosomal protein L18 | ADP39349.1 | HMPREF0421_21267 | Gardnerella vaginalis 14019 |
| ileS | isoleucine--tRNA ligase | ADP38567.1 | HMPREF0421_20485 | Gardnerella vaginalis 14019 |
| rpsD | ribosomal protein S4 | ADP39181.1 | HMPREF0421_21099 | Gardnerella vaginalis 14019 |
| valS | valine--tRNA ligase | ADP39401.1 | HMPREF0421_21319 | Gardnerella vaginalis 14019 |
| gcp | putative glycoprotease GCP | ADP38822.1 | HMPREF0421_20740 | Gardnerella vaginalis 14019 |
| ffh | signal recognition particle protein | ADP39221.1 | HMPREF0421_21139 | Gardnerella vaginalis 14019 |
| ftsY | signal recognition particle-docking protein FtsY | ADP38181.1 | HMPREF0421_20095 | Gardnerella vaginalis 14019 |

**Table S2.**
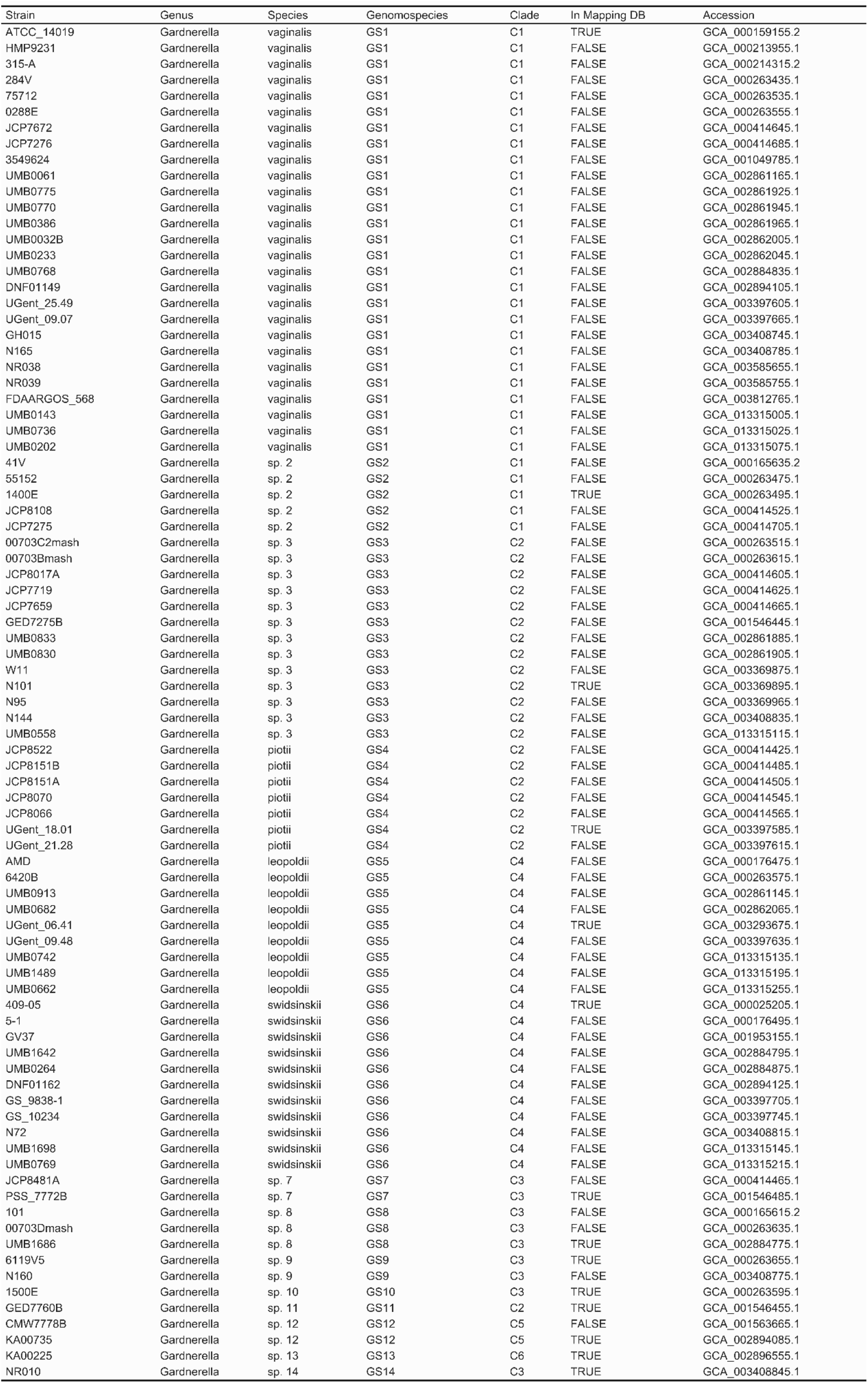
Reference Gardnerella Genomes

**Table S3.**
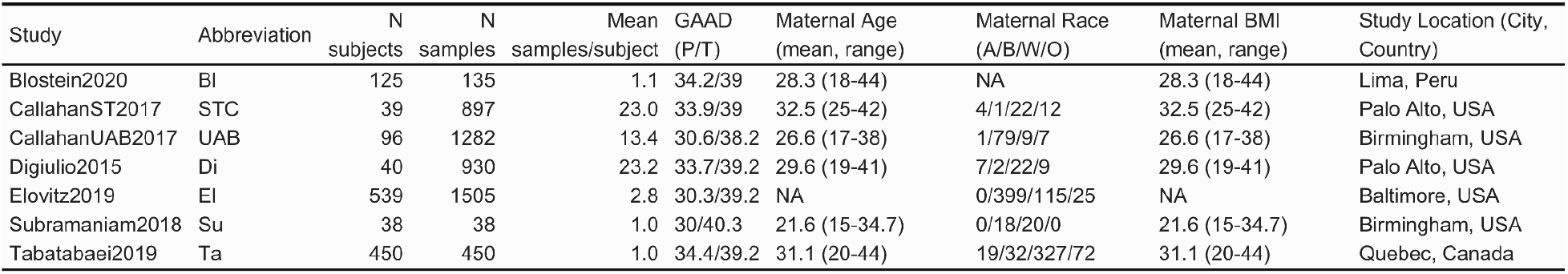
V4 16S Amplicon Studies

## Supplementary Figures

**Figure S1.**
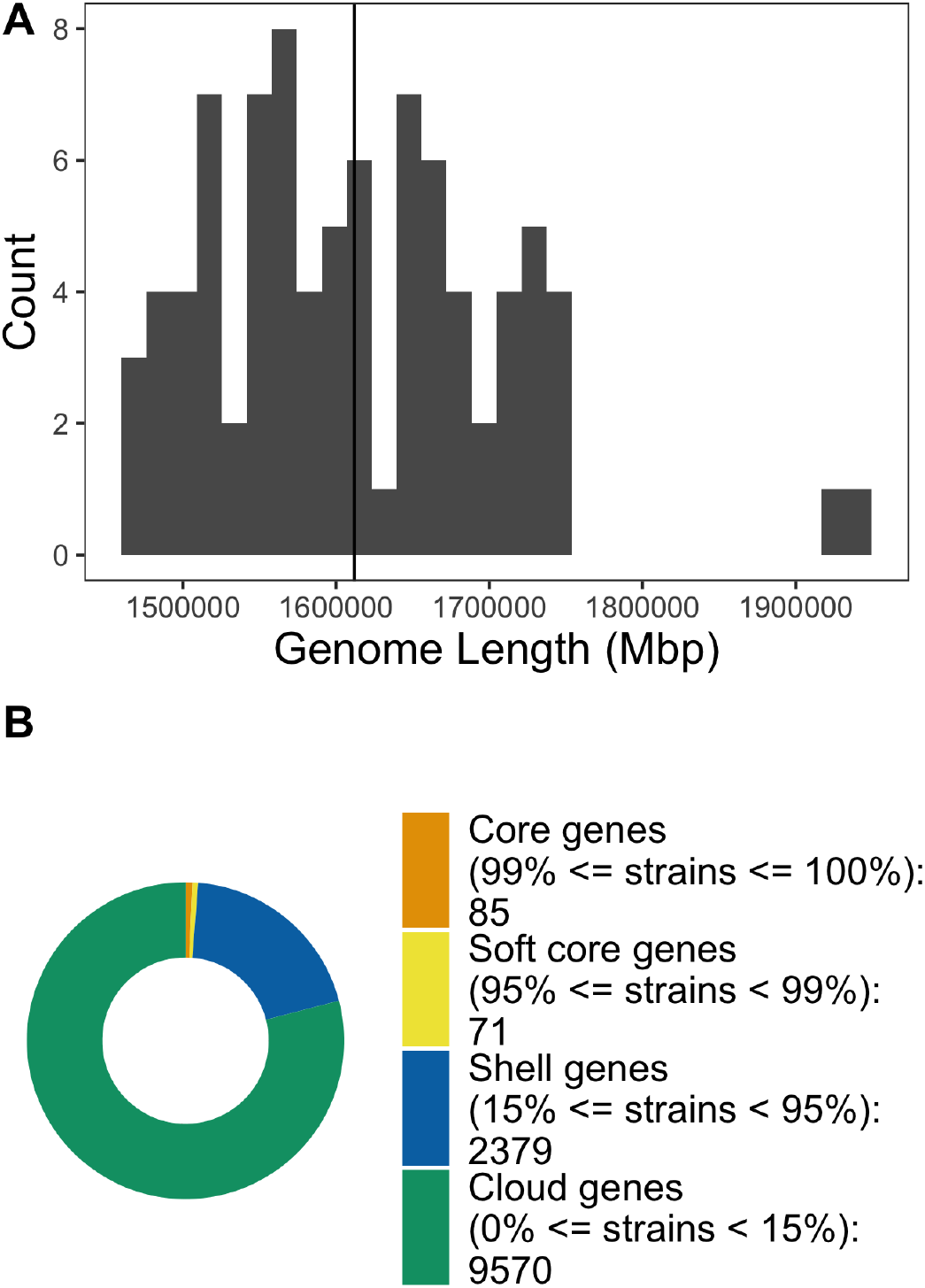
A) Full length of genomes used in reconstructing phylogeny. Vertical line represents mean genome size (1.6Mbp). B) Core, soft core, shell, and cloud genes in the total pangenome.

**Figure S2.**
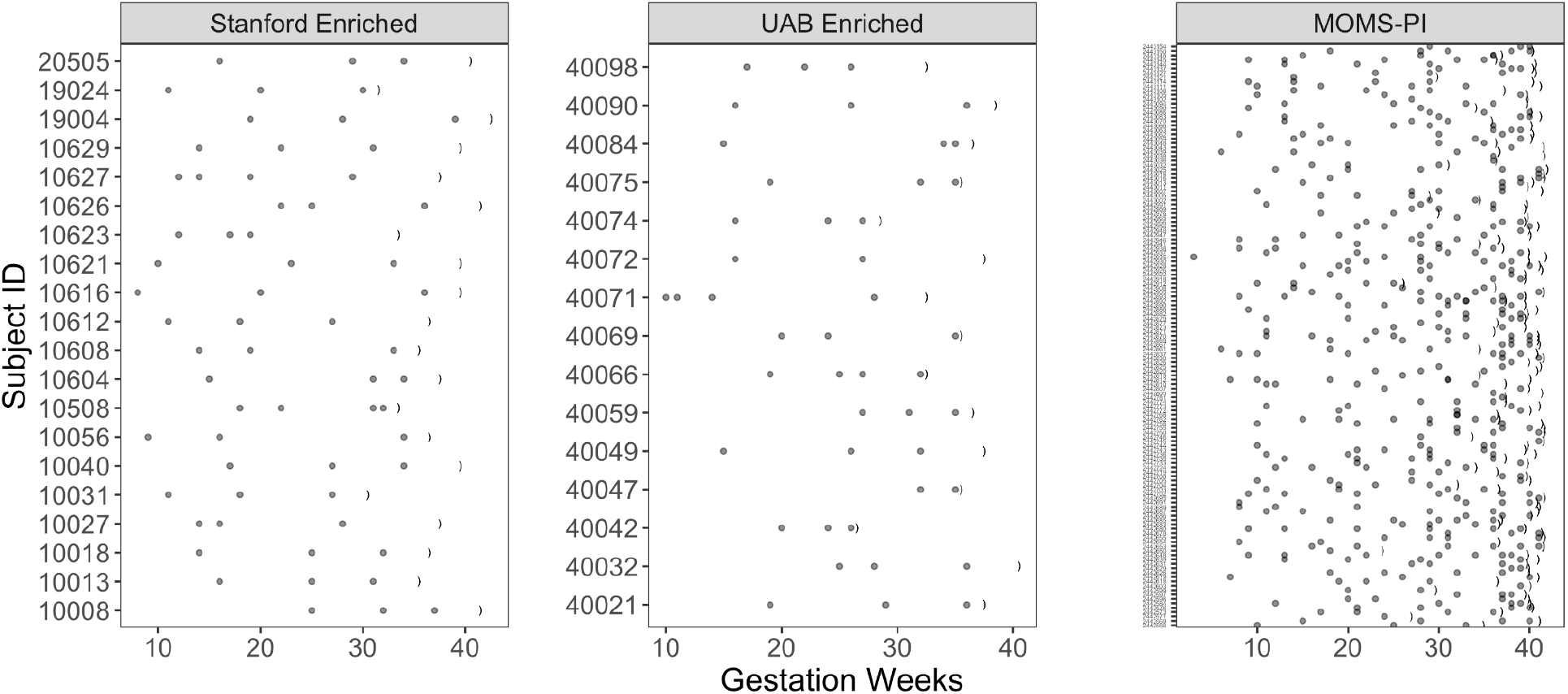
Sampling schedule in the Stanford, UAB, and MOMS-PI cohorts. Points show vaginal swab samples, and parentheses indicate the gestational week of delivery. The median week of sampling in the Stanford cohort was 22, and the interquartile range (IQR) was 16–31 wk. The median week of sampling in the UAB cohort was 26, and the IQR was 19–32 wk. The median week of sampling in the MOMS-PI cohort was 29 weeks, and the IQR was 20-36 weeks.

**Figure S3.**
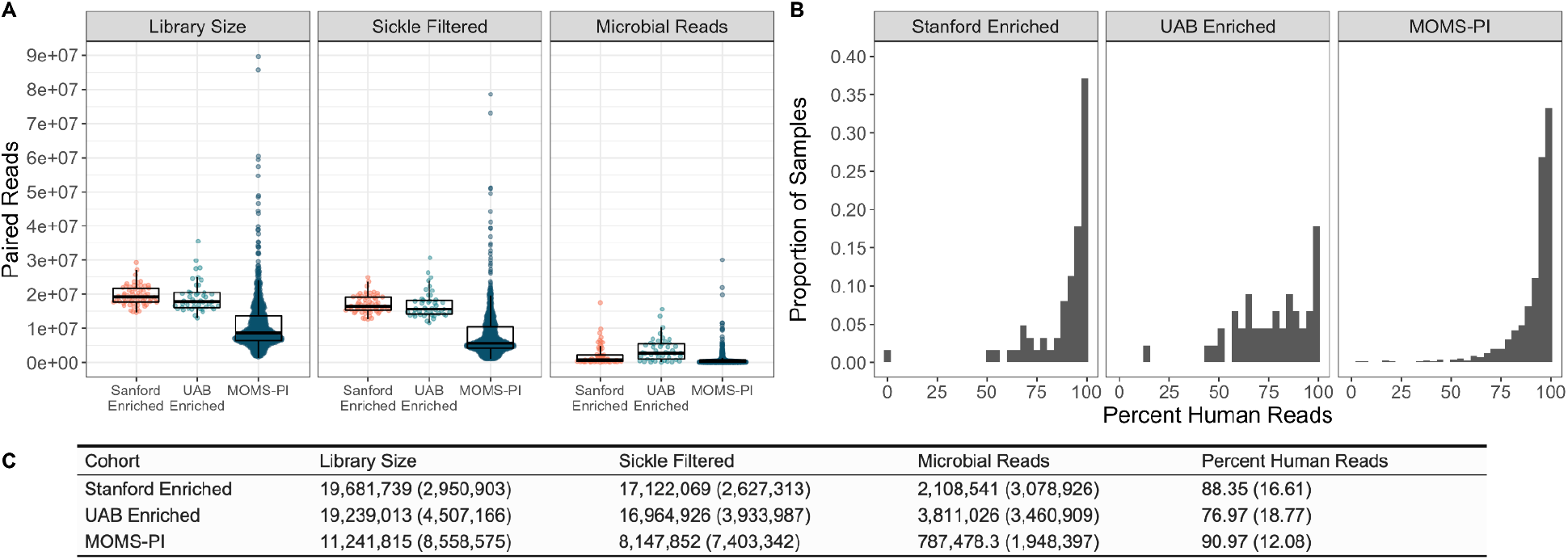
Library Sizes and Sequence Filtering. A) Library sizes and counts of paired reads after filtering with Sickle for quality and for human reads in the Stanford Enriched, UAB Enriched, and MOMS-PI cohorts. B) Percent of human reads in each sample. C) Mean (and standard deviation) of library size, and filtered reads after sickle and human filtering, and percent of human reads in samples.

**Figure S4.**
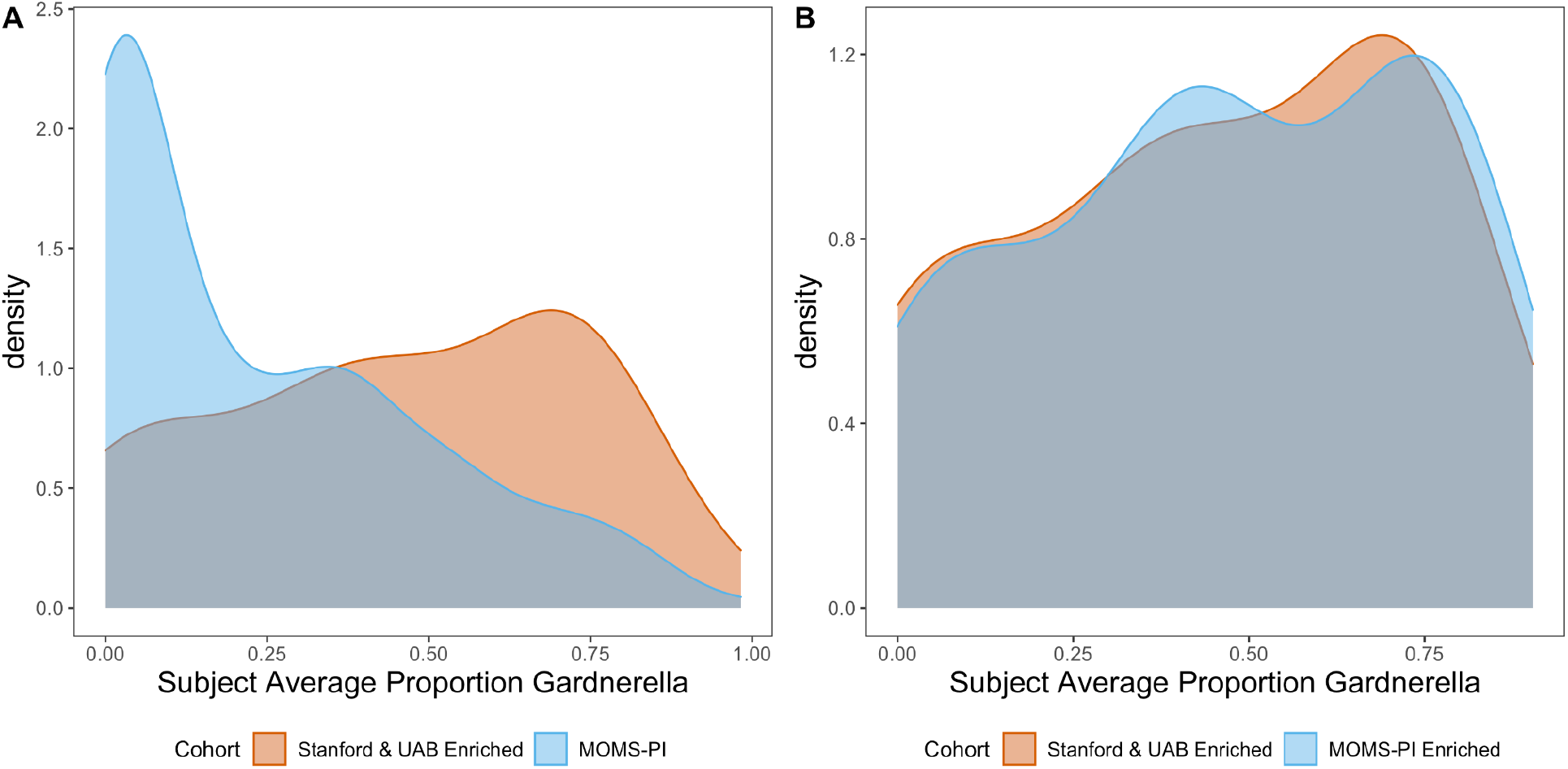
MOMS-PI Cohort Enriched for subject abundance of Gardnerella. Comparison of subject average abundance of *Gardnerella* in A) full MOMS-PI cohort and B) subset enriched for *Gardnerella* to Stanford and UAB cohorts which were selected for shotgun metagenomic sequencing based on *Gardnerella* abundance in subjects as determined by 16S rRNA gene amplicon sequencing.

**Figure S5.**
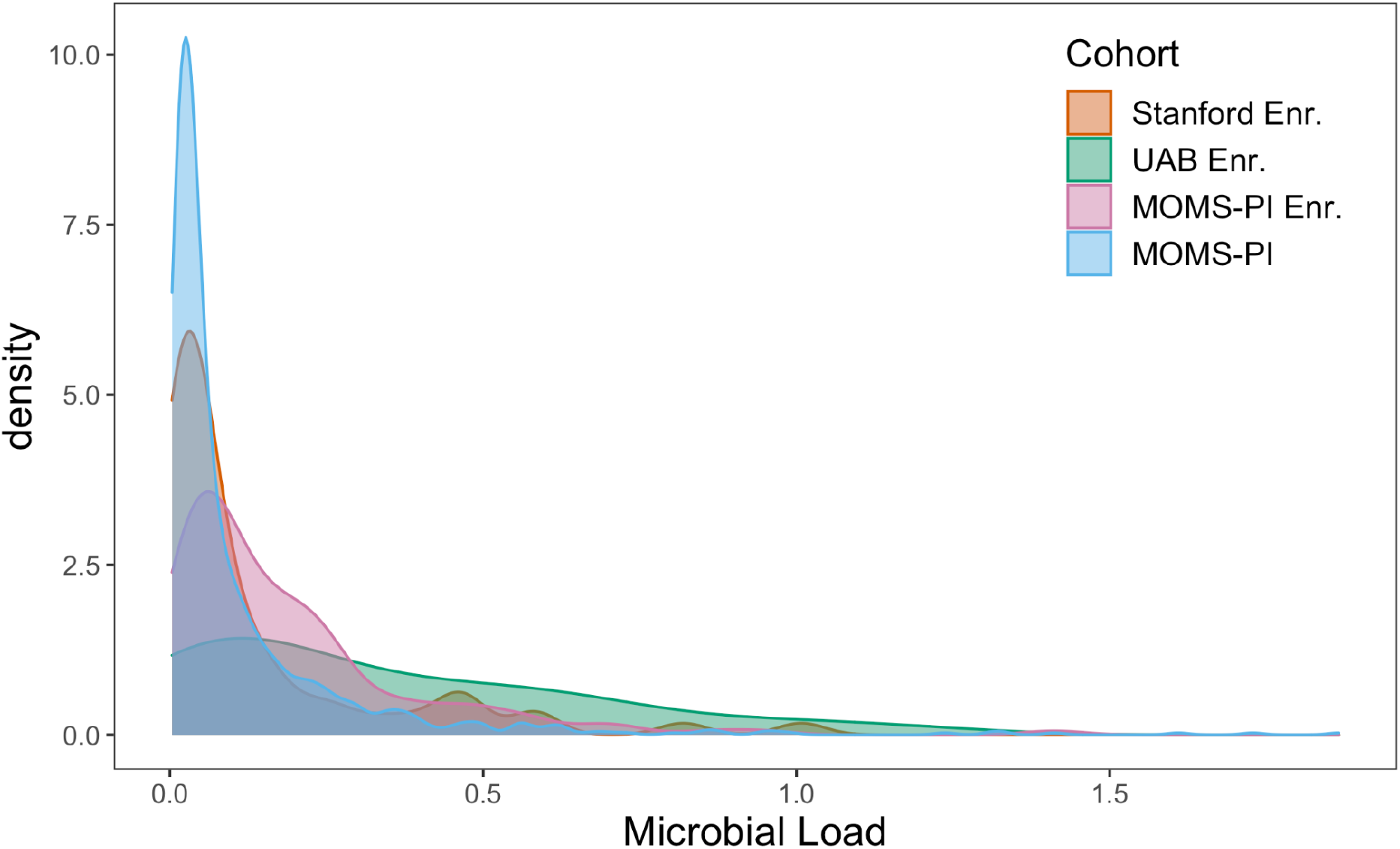
Density plot of the bacterial load (ratio of bacterial reads to human reads) of samples in the four cohorts.

**Figure S6.**
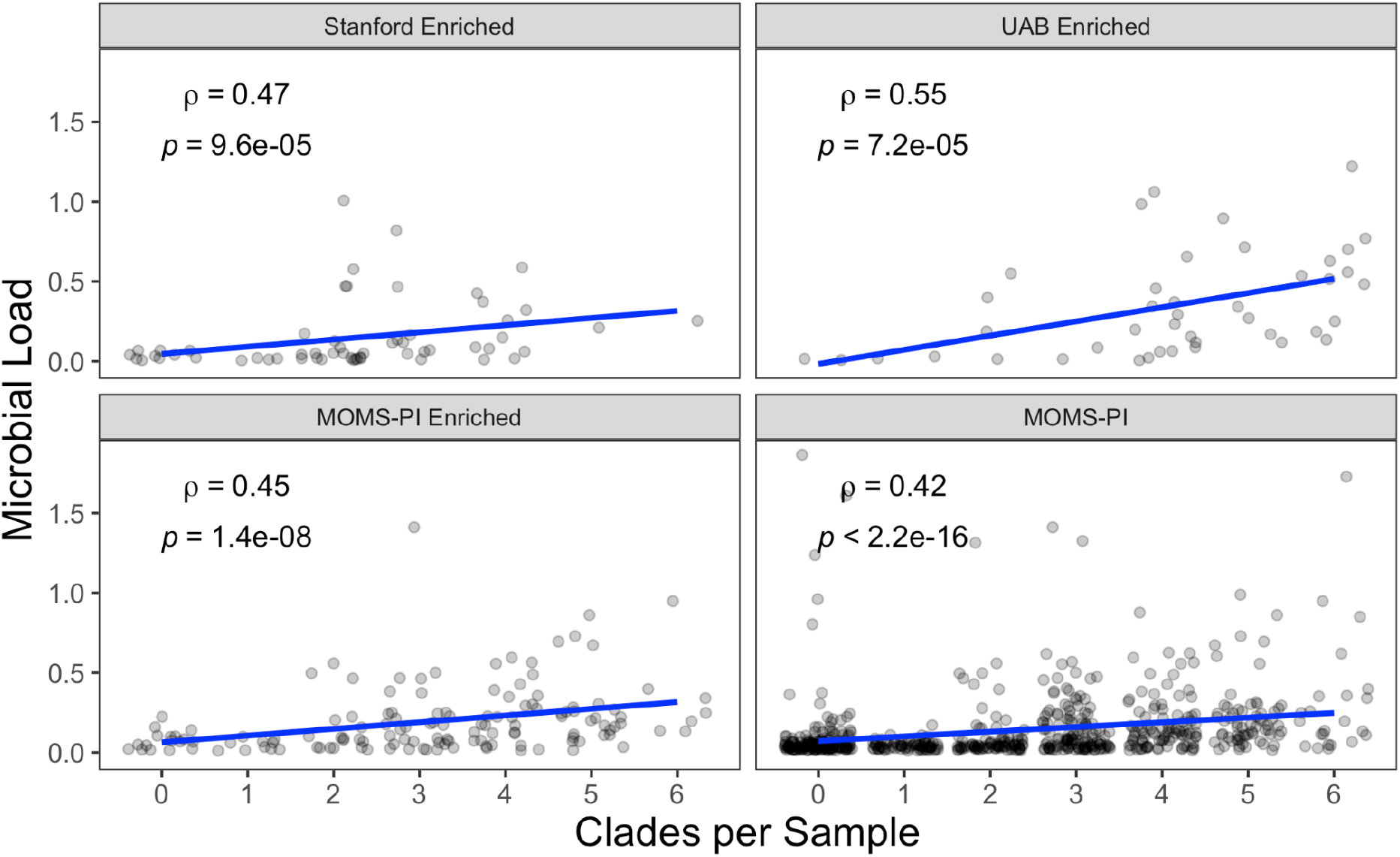
Bacterial load vs. clades per sample after rarefying. Human-read-filtered samples were rarefied to a common depth of 100,000 reads and clades per sample were compared to microbial load using Spearman’s rank correlation. Bacterial load was calculated from un-rarefied reads.

**Figure S7.**
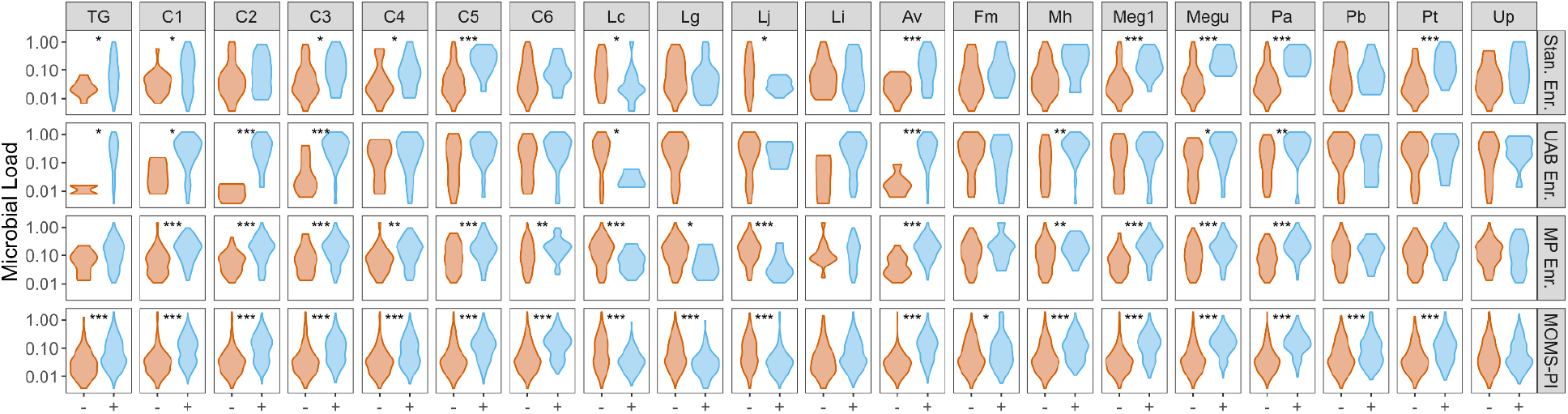
Bacterial load by presence-absence of *Gardnerella* clades and species in the vaginal microbiome. *p* values reflect one-directional Wilcoxon Rank Sum tests. *: *p* < 0.05; **: *p* < 0.01; ***: *p* < 0.001. Direction of Wilcoxon Rank Sum test determined by previous associations with microbial load: *Gardnerella* and other anaerobes associated with increased microbial load and *Lactobacillus* spp. associated with decreased microbial load. Abbreviations: TG = Total *Gardnerella*, Lc = *Lactobacillus crispatus*, Av = *Atopobium vaginae*, Fm = *Finegoldia magna*, Mh = *Mycoplasma hominis*, Meg1 = *Megasphaera* genomospecies type 1, Megu = *Megasphera* unclassified, Pa = *Prevotella amni*, Pb = *Prevotella bivia*, Pt = *Prevotella timonensis*, Up = *Ureaplasma parvum*

**Figure S8.**
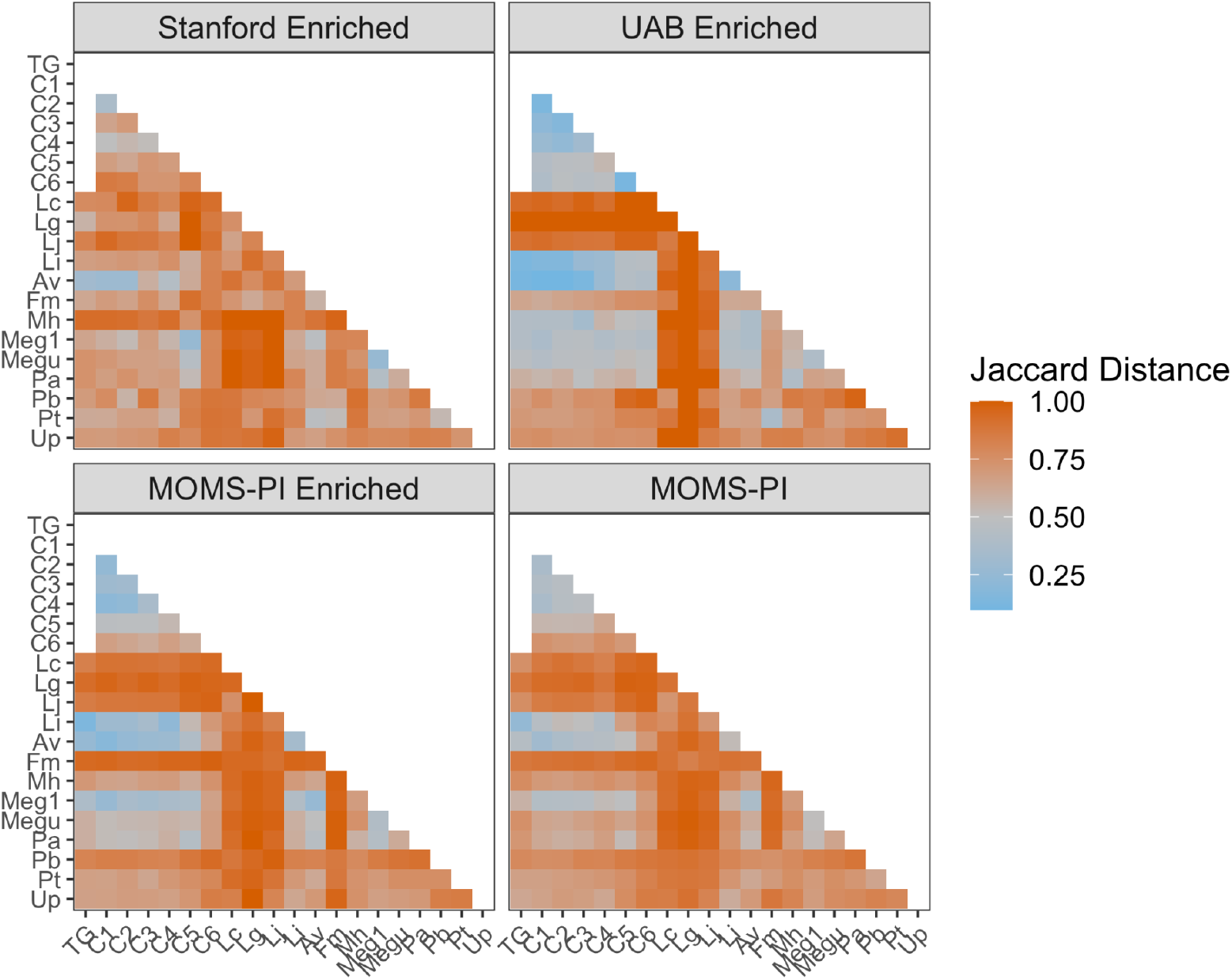
Jaccard distances among *Gardnerella* clades and other key taxa in the vaginal microbiome. Abbreviations: TG = Total *Gardnerella*, Lc = *Lactobacillus crispatus*, Av = *Atopobium vaginae*, Fm = *Finegoldia magna*, Mh = *Mycoplasma hominis*, Meg1 = *Megasphaera* genomospecies type 1, Megu = *Megasphera* unclassified, Pa = *Prevotella amni*, Pb = *Prevotella bivia*, Pt = *Prevotella timonensis*, Up = *Ureaplasma parvum*

**Figure S9.**
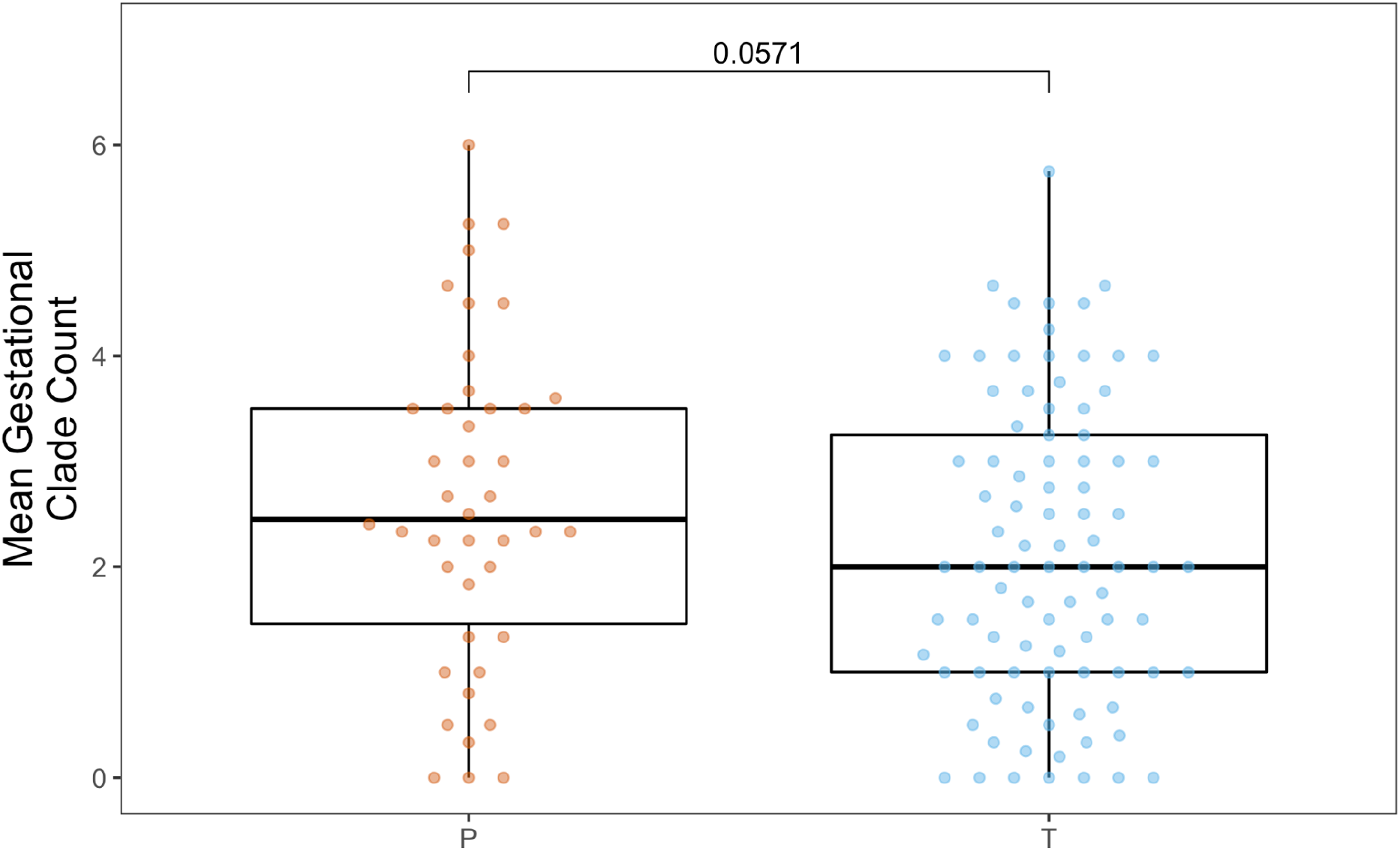
Mean gestational clade counts versus PTB after rarefying. Human-read-filtered samples were rarefied to a common depth of 100,000 reads and mean clades per sample per subject were compared among subjects who delivered at term and preterm by a Wilcoxon Rank Sum test. Clades were considered present in a sample if the relative abundance was greater than 0.1%.

**Figure S10.**
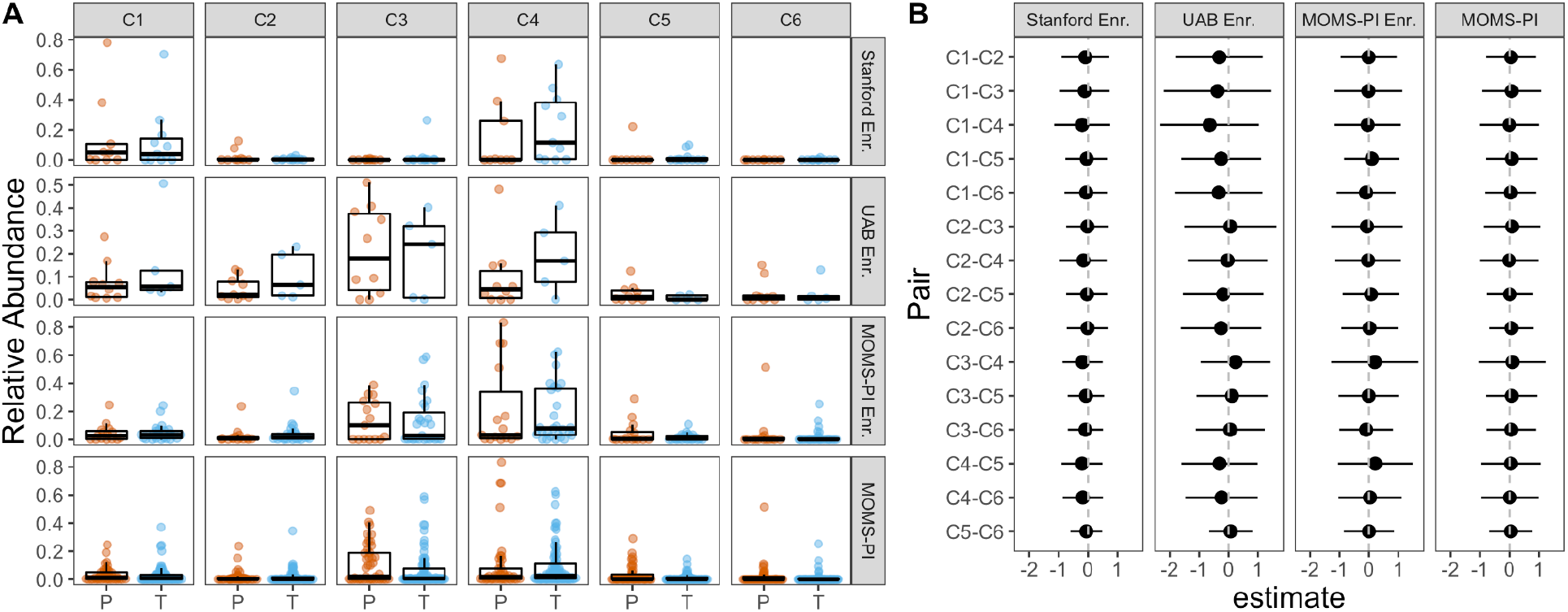
Comparisons among relationships between clade abundances and preterm birth. A) Mean gestational relative clade abundance by preterm birth status. B) Estimates of logistic regression coefficients +/− standard deviations comparing pairwise differences in relative abundance in term and preterm births. No coefficients were found to be significant suggesting that associations among clades with preterm birth did not vary.

**Figure S11.**
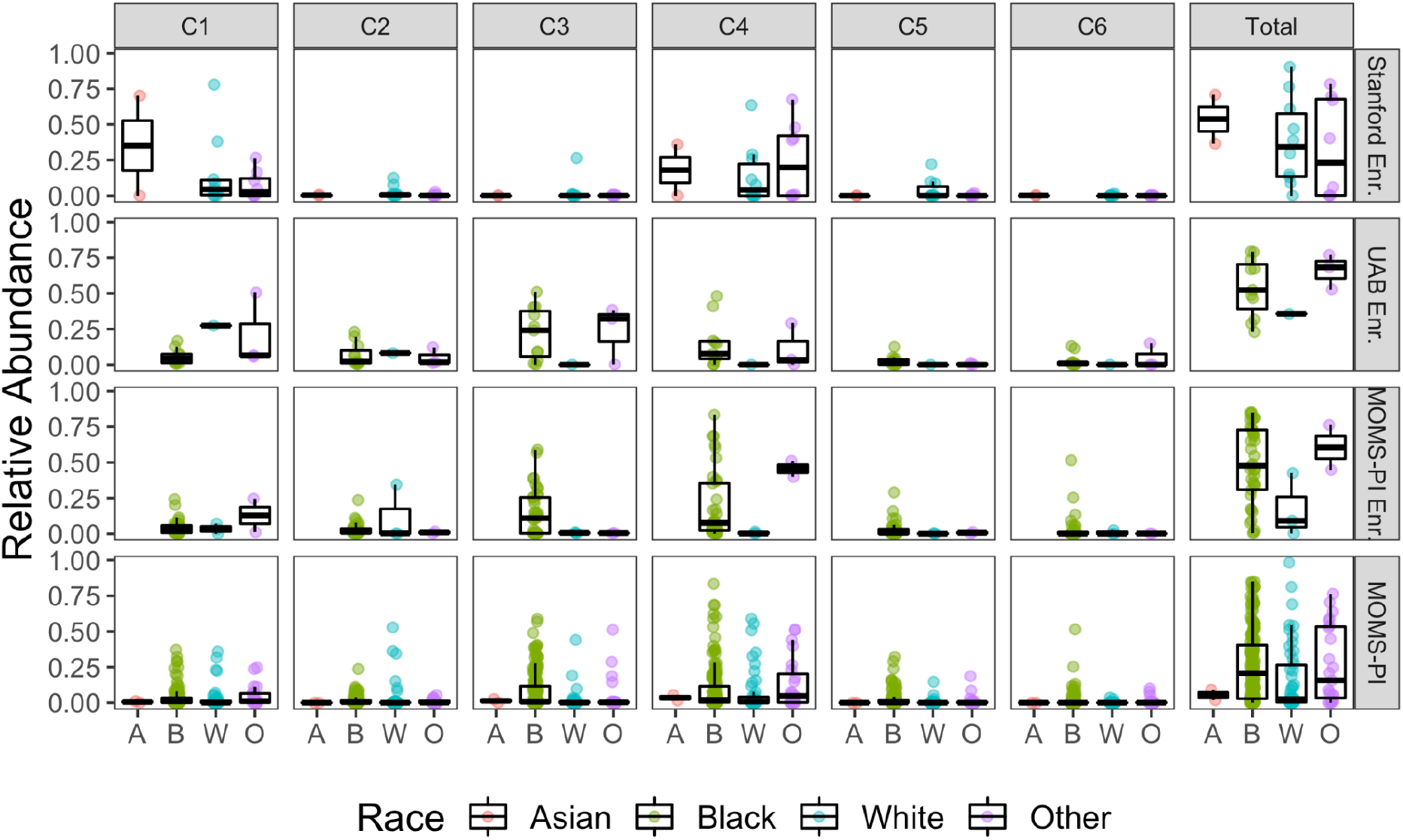
Subject average relative abundance of total *Gardnerella* and specific clades by subject race.

**Figure S12.**
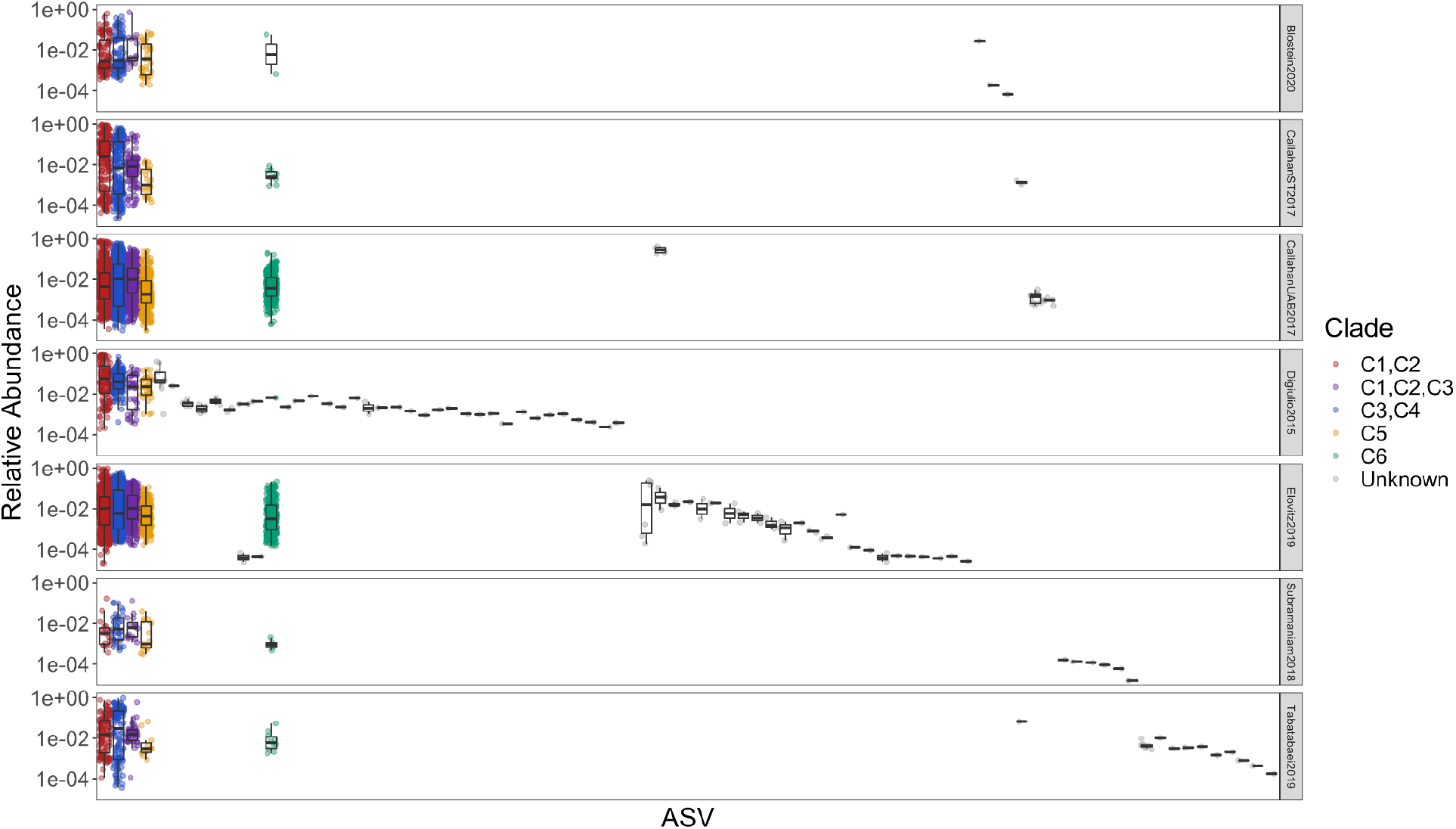
Relative abundance of all *Gardnerella* ASVs across all studies. Note log_10_ y-axis for easier viewing of ASVs with low abundance.

